# Combined Pan-, Population-, and Phylo-Genomic Analysis of *Aspergillus fumigatus* Reveals Population Structure and Lineage-Specific Diversity

**DOI:** 10.1101/2021.12.12.472145

**Authors:** Lotus A. Lofgren, Brandon S. Ross, Robert A. Cramer, Jason E. Stajich

## Abstract

*Aspergillus fumigatus* is a deadly agent of human fungal disease, where virulence heterogeneity is thought to be at least partially structured by genetic variation between strains. While population genomic analyses based on reference genome alignments offer valuable insights into how gene variants are distributed across populations, these approaches fail to capture intraspecific variation in genes absent from the reference genome. Pan-genomic analyses based on *de novo* assemblies offer a promising alternative to reference-based genomics, with the potential to address the full genetic repertoire of a species. Here, we use a combination of population genomics, phylogenomics, and pan-genomics to assess population structure and recombination frequency, phylogenetically structured gene presence-absence variation, evidence for metabolic specificity, and the distribution of putative antifungal resistance genes in *A. fumigatus*. We provide evidence for three primary populations of *A. fumigatus*, structured by both gene variation (SNPs and indels) and distinct gene presence-absence variation with unique suites of accessory genes present exclusively in each clade. Accessory genes displayed functional enrichment for nitrogen and carbohydrate metabolism, suggesting that populations may be stratified by environmental niche specialization. Similarly, the distribution of antifungal resistance genes and resistance alleles were often structured by phylogeny. *A. fumigatus* demonstrated exceptionally high levels of recombination and one of the largest fungal pan-genomes reported to date including many genes unrepresented in the Af293 reference genome. These results highlight the inadequacy of relying on a single-reference genome-based approach for evaluating intraspecific variation, and the power of combined genomic approaches to elucidate population structure, genetic diversity, and putative ecological drivers of clinically relevant fungi.

## INTRODUCTION

*Aspergillus fumigatus* is one of the most common etiological agents of human fungal disease (Tekaia & Latgé, 2005; Department of Health and Human Services. CDC., 2019). The spectrum of diseases attributed to *A. fumigatus* is remarkable. In immunocompromised patients, invasive *A. fumigatus* infection causes up to 90% mortality, even with aggressive treatment (Latgé & Chamilos, 2019), an outcome that is further complicated by the increasing presence of triazole resistant strains (Howard *et al*., 2006). Phenotypic heterogeneity in growth and virulence is well documented in *A. fumigatus* (Kowalski *et al*., 2016, 2019; Keller, 2017; Caffrey-Carr *et al*., 2017; Ries *et al*., 2019) and thought to be partially structured by genetic variation between strains (Rokas *et al*., 2020). These intraspecific genetic differences likely represent both gene variants (insertions/deletions and single nucleotide polymorphisms) (Ballard *et al*., 2018), and differences in gene presence-absence, copy number, and structural arrangements (Zhao & Gibbons, 2018; Steenwyk *et al*., 2020). While population-genomic analyses based on reference genome alignment have provided valuable insights into how gene variants are distributed across populations, these approaches fail to capture intraspecific variation in genomic regions absent from the reference genome. In contrast to reference-based genomics, pan-genomic analyses based on *de novo* assemblies offer the potential to address the full genetic repertoire of a species. For *A. fumigatus* pathogenesis and virulence, identification of strain specific genes and alleles is expected to further clarify the role of fungal genetic variation in disease outcomes.

The concept of a pan-genome, here defined as all genetic elements present across a species (Tettelin *et al*., 2005), recognizes that while many genes are fixed within a population and present in all individuals (core genes) others display substantial presence-absence variation (dispensable or accessory genes). Core genes are expected to be enriched in housekeeping functions, and clustered in protected areas of chromosomes, where they are subject to low rates of recombination and mutation (Croll & McDonald, 2012). Conversely, accessory genes are more likely to encode lineage specific proteins that facilitate environmental adaptation (Medini *et al*., 2005), such as genes involved in secondary metabolism and niche specificity, and tend to be concentrated in highly variable, rapidly evolving sub-telomeric regions and adjacent to Transposable Elements (TEs) (Fedorova *et al*., 2008; Brown *et al*., 2010; Faino *et al*., 2016; Wyatt *et al*., 2020). Variation in the accessory genome is essential to facilitating diversifying selection and adaptation to population-specific environmental pressures, and may be particularly important for fungal pathogen adaptation (Croll & McDonald, 2012; Badet & Croll, 2020).

In ecological settings, *A. fumigatus* is a ubiquitous plant litter saprophyte, encoding a wide range of carbohydrate-active enzymes involved in the decay of organic matter, and playing important roles in carbon and nutrient cycling (Tekaia & Latgé, 2005; Willger *et al*., 2009). *A. fumigatus* reproduces primarily asexually, via the production of prolific, stress-resistant, hydrophobic conidia (Dagenais & Keller, 2009). This combination of traits contribute to the high dispersibility of the species, where conidia are rendered airborne by the slightest wind currents, or easily caried to new locations by water, swarming soil bacteria, and soil invertebrates (Kwon-Chung & Sugui, 2013). Due to this exceptional dispersibility and the assumption of extreme substrate generalism, *A. fumigatus* was originally thought to represent a single homogenous population (Pringle *et al*., 2005; Rydholm *et al*., 2006). However, recent investigations into *A. fumigatus* population structure have found mixed evidence for population stratification, with results heavily dependent on the methods used for analysis, and the number of isolates under consideration (Abdolrasouli *et al*., 2015; Ashu *et al*., 2017; Sewell *et al*., 2019; Barber *et al*., 2021). Interestingly, previous studies have shown little to no correlation of population structure with geography (Klaassen *et al*., 2012; Sewell *et al*., 2019), with genetically identical (clonal) isolates collected from disparate locations across the globe (Sewell *et al*., 2019). This lack of geographic structure would seem to support a model of panmixia, but geography is not the only potential force structuring fungal populations, which must adapt to a plethora of environmental stressors and niche opportunities. However, niche specificity, and the potential for environmental drivers to structure population stratification in *A. fumigatus* have yet to be investigated.

Because *A. fumigatus* is thought to reproduce primarily clonally (and rarely sexually) (O’Gorman *et al*., 2009), population structure is likely to be complicated by local clonal population bursts followed by dispersal. One well studied example of this is the movement of *cyp51A* resistance alleles across the world. The *cyp51A* gene encodes 14 alpha sterol demethylase and is the primary drug target for azole antifungals (Hagiwara *et al*., 2016). Environmental pressures for allelic variation at this critical azole target are hypothesized to come primarily from the wide-spread use of agricultural antifungals (Chowdhary *et al*., 2013). Human-to-human transmission was originally thought to represent a dead-end for the species, however recent evidence support occasional hospital-related transmission (Lemaire *et al*., 2018; Engel *et al*., 2019), and the possibility for clinical antifungal treatment to represent a secondary source of antifungal pressure. Although the *cyp51A* mutation has been found in multiple genetic backgrounds (Abdolrasouli *et al*., 2015), the distribution of drug resistant isolates across the phylogeny is non-random, with resistant strains showing low levels of genetic diversity and close genetic relatedness, likely representing high levels of clonal reproduction and a selective sweep for resistant phenotypes (Klaassen *et al*., 2012; Sewell *et al*., 2019). Although *cyp51A* mutations are the best studied drug resistance mechanism in *A. fumigatus*, mutations in *cyp51A* only account for an estimated 43% of resistant isolates (Bueid *et al*., 2010). Although many other genes have been implicated in Azole resistance (Chen *et al*., 2020), it is unknown whether mutations in these genes are similarly structured by phylogeny.

Whereas widely distributed generalist species with large effective population sizes are assumed to have larger pan-genomes, frequent clonal reproduction is thought to limit pan-genome size (McInerney *et al*., 2017; Badet & Croll, 2020). Generally, the evolution of large pan-genomes implies frequent gene flow, which may take the form of sexual or pseudo-sexual exchange, or significant horizontal gene transfer (Naranjo-Ortiz & Gabaldón, 2020). In open pan-gnomes, the ratio of core/accessory genes is expected to decrease with an increasing number of genomes analyzed. Conversely, closed pan-genomes will quickly reach a saturation point at which adding new genomes to the analysis adds few new genes to the total pool. It is unclear how a species a like *A. fumigatus*, which demonstrate ubiquitous geographic distribution and assumed substrate generalism, primary clonal reproduction, but potentially high recombination rates (Auxier *et al*., 2022), fits into these expectations.

## METHODS

### DNA preparation

All strains sequenced in this project were isolated on Aspergillus Minimal Medium (Cove, 1966) by picking single germinated spores after 16 hrs of growth at 30 °C. Cultures for DNA extractions were grown in liquid minimal media with 1% (w/v) D-Glucose, 0.5% Yeast Extract (w/v, Beckton-Dickinson), 20 ml 50x Salt Solution, 1 ml Trace Elements solution, 20 mM NaNO_3_, pH adjusted to 6.5 using NaOH, and autoclaved for 20 minutes at 121 °C.

### Genome Sequencing

After 24 hours of growth, cultures were lyophilized for ∼10 hours, and homogenized using a bead beater (1 min with 2.3 mm beads). DNA was extracted using a LETS buffer protocol (Griffiths *et al*., 2006) modified with the addition of a one hour RNAse treatment prior to phenol-chloroform extraction. DNA concentration was quantified using a Qubit 2.0 Fluorometer (Invitrogen) with the Broad Range protocol. Genomic sequencing was caried out on either Illumina NovaSeq 6000 or NextSeq 500 machines. DNA sequencing libraries were prepared using either the NEBNext Ultra II DNA Library Prep Kit (for NovaSeq 6000 sequenced genomes), or the SeqOnce DNA library kit utilizing Covaris mechanized shearing (for the NextSeq 500 sequenced isolates), both following manufacturer recommendations with paired end library construction and barcoding for multiplexing. All genome sequencing data generated for this project was deposited into the NCBI Sequence Read Archive under BioProject no. PRJNA666940. In addition to the 62 newly sequenced strains, a large library of strains (∼220) previously published by our lab and others were downloaded from NCBI’s SRA and assembled and annotated.

### De-Novo assembly and annotation

Genomes were assembled *de novo* with the Automatic Assembly For The Fungi (AAFTF) (https://github.com/stajichlab/AAFTF, DOI: 10.5281/zenodo.1620526.) The AAFTF *filter* step trim and quality filter reads with BBmap (https://sourceforge.net/projects/bbmap) and the AAFTF *assemble* step was used to assemble the reads with SPAdes (Bankevich *et al*., 2012). Resulting contigs were screened for bacterial contamination using the AAFTF *sourpurge* step which relies on sourmash (Titus Brown & Irber, 2016) searching a database of Genbank microbial sequence sketches (v.lca-mark2; https://osf.io/vk4fa/). Duplicates were removed with AAFTF *rmdup* by aligning contigs to themselves with minimap2 (Li, 2018), and contigs were further polished for accuracy using the AAFTF *polish* step which relies on Pilon (Walker *et al*., 2014). Scaffolds from contigs were inferred by aligning to the reference Af293 genome and scaffolding with ragtag v 1.0.0 (Alonge *et al*., 2021). To assess contig and scaffold genome completeness BUSCO v4.0.5 (Simão *et al*., 2015) was run on all genomes and only isolates with >95% total coverage were used in the analysis. Read depth was assessed using BBTools (https://jgi.doe.gov/data-and-tools/bbtools/) and only assemblies with an average read depth >10X coverage were retained. This resulted in a total of 260 quality-filtered strains, including 62 new strains and 198 strains previously published strains (Table S1), including the re-sequenced reference strain AF293, which was included as a control for variant analysis conducted relative to the Af293 v.55 curated reference downloaded from FungiDB.

Repeat regions were masked by funannotate *mask* (https://github.com/nextgenusfs/funannotate<u>, DOI:</u> <u>10.5281/zenodo.1134477</u>) using RepeatMasker (Tarailo-Graovac & Chen, 2009) with repeats built using RepeatModler (Flynn *et al*., 2020) compiled into a custom library (available in this the project’s GitHub) plus those fungal repeat families curated in RepBase v. 20170127 (Bao *et al*., 2015). The *train* command in funannotate was used to run Trinity (Grabherr *et al*., 2011) for transcript assembly and alignments to generate highly polished gene models based on splice-site aware alignment of the assembled RNA-Seq of *A. fumigatus* growth on sugarcane bagasse (PRJNA376829) (de Gouvêa *et al*., 2018). Models were further refined by PASA and the best set was chosen for input in training gene predictors (Haas *et al*., 2003). Gene prediction was carried out using funannotate *predict* running Augustus (Stanke & Morgenstern, 2005) and SNAP (Korf, 2004) by first training on the evidenced based training models from the *train* step. Gene prediction was then run with these trained parameters using exon evidence based on RNAseq and protein alignments generated by DIAMOND v.2.0.2 (Buchfink *et al*., 2015) and polished with Exonerate v.2.4.0 (Slater & Birney, 2005) on the RNA-seq and Swissprot proteins (UniProt Consortium, 2019). Additional *ab initio* models were predicted using GeneMark v.4.59 (Ter-Hovhannisyan *et al*., 2008), GlimmerHMM v.3.0.4 (Majoros *et al*., 2004), and CodingQuarry v.2.0 (Testa *et al*., 2015). Evidence for consensus gene models were produced with EVidenceModeler v.1.1.1 (Haas *et al*., 2008). The tool funannotate *annotate* was run to annotate protein domains and make functional predictions using sequence similarity with the databases InterProScan v.5.45-80.0 (Finn *et al*., 2017), eggNOG v.1.0.3 (Huerta-Cepas *et al*., 2017) dbCAN2 v.9.0 (Zhang *et al*., 2018), UniprotDB v.2020_04, antiSMASH v.5.1.2 (Blin *et al*., 2019) and MEROPS v.12.0 (Rawlings *et al*., 2018).

### Pan-genomics

We used two methods to determine pan-genome gene family clusters: OrthoFinder v2.5.2 (Emms & Kelly, 2018) and Pangenome Iterative Refinement and Threshold Evaluation (PIRATE) v1.0.4 (Bayliss *et al*., 2019). OrthoFinder was run on protein fastA files with -op option to run similarity searches as parallel jobs on the HPC, and the -S diamond_ultra_sens option for sequence search using DIAMOND. PIRATE was run on nucleotide fastA files, with parameters: -s “85,86,87,88,89,90,91,92,93,94,95,96,97,98,99,100” -k “--cd-low 100 -e 1E-9 --hsp-prop 0.5” -a -r. Pan-genome clustering results were analyzed in the R programing environment using custom scripts (all scripts are available at the DOI listed in the data availability section). For our analysis ‘core’ genes were defined as present in >95% of the strains (n > 247), ‘accessory genes’ were defined as present in more than one, and less than 95% of the strains (n >1 and <=247) and singletons were defined as present in only a single isolate. Significant differences in the abundance of accessory and singleton gene families per clade were assessed using a permutation test implemented in the R package *perm* (Fay & Shaw, 2010) over 9999 permutations, to account for violation of variance assumptions. Gene family accumulation curves were calculated using the specaccum() function over 100 iterations in R package *vegan* v2.5.7 (Dixon, 2003). Analysis of clade-specific gene family absence was defined as a gene family absent in all isolates of that clade, but present in both of the other clades and in >90% of the isolates from at least one of those clades. Additional screening for clade-defining gene family absences was also conducted, defined as absent in all isolates of that clade, but present in >95 percent of all isolates from both of the other clades. Secreted proteins were predicted using the programs Signal P5 (Armenteros *et al*., 2019) and Phobius (Käll *et al*., 2004). Phobius predictions were further refined by both the presence of a signal peptide, and the absence of transmembrane domains. Significant differences in proportions of secreted gene families relative to all gene families were assessed using pairwise proportion tests with Bonferroni adjustment for multiple comparisons using function pairwise.prop.test() in the R *stats* package at *p* < 0.05.

Gene Ontology (GO) enrichment analysis was performed by assigning Interpro and GO annotations to the longest representative of each gene family clustered in OrthoFinder. GO enrichment analysis was then preformed for each clade, on gene all families unique to that clade using the R package *topGO* v.2.42.0 with the functions new() and runTest() with parameters weight01 and nodeSize=6 (DOI: 10.18129/B9.bioc.topGO). Significant enrichment was assessed for each gene count category (core, accessory, singleton) and for clade-specific gene families relative to the set of all InterPro annotations with associated GO terms in the *A. fumigatus* pan-genome (n=32,857), using a Fishers Exact test at *p* < 0.05. CAZyme (Carbohydrate-Active enZyme) annotations were assigned in funannotate which relies on hmmsearch to search the dbCAN database for HMM profiles (Zhang *et al*., 2018). Homogeneity of variance assumptions were checked using the leveneTest() function from the *car* package v.3.0.11 and normalcy checked using the shapiro.test() function from the *stats* package v.4.0.1 in R. Because variance and normalcy assumptions could not be met, non-parametric Kruskal Wallace tests were employed using the kruskal.test() function in the R *stats* package v.4.0.1 with Bonferroni adjustment for multiple comparisons using the p.adjust() function in R, and considered for further evaluation at *p* < 0.001. Post-hoc comparisons using Dunn’s test were implemented using the dunnTest() function with Bonferroni adjustment for multiple comparisons in the R package *FSA* v.0.9.0. CAZymes significantly different between the three clades were normalized per CAZyme family on a 0-1 scale for visualization and mapped onto the phylogeny using the gheatmap() function in the R package *ggtree* v.3.1.2. To validate our OrthoFinder clustered gene families, we represented each gene family with the longest sequence, and used BLASTP v. 2.12.0 (e-val < 1e^-15^) to match gene families to their respective genes annotated in the Af293 reference genome. To further investigate the presence-absence distribution of accessory genes with a role in nitrogen, carbohydrate, and phosphorus metabolism, we extracted all Af293 genes from FungiDB with GO annotations that matched organonitrogen compound metabolic process (GO: 1901564, n=1,775 genes), carbohydrate metabolic process (GO: 0005975, n=467 genes), or organophosphate biosynthetic process (GO: 0090407, n=215 genes). We then examined gene families with orthologues in Af293, to test for presence-absence variation in these genes between the three clades. Significant differences in gene abundance between the three clades was evaluated using the same strategy outlined above to evaluate CAZyme abundance.

### Population genomics

The Af293 reference genome was downloaded from FungiDB (v.46) (Basenko *et al*., 2018). Sequence reads for each strain were aligned to Af293 using BWA v0.7.17 (Li & Durbin, 2009). Using samtools v1.10 (Li *et al*., 2009), the fixmate and sort commands were run and files were converted to the BAM format. Duplicate reads were removed, and reads were flagged using MarkDuplicates and indexed with Build BamIndex in picard tools v.2.18.3 (http://broadinstitute.github.io/picard). Reads that overlapped INDELS were realigned using RealignerTargetCreator and IndelRealigner in the Genome Analysis Toolkit GATK v.3.7 (Auwera *et al*., 2013). Variants were called relative to Af293 using HaplotypeCaller in GATK v.4.0, with filtering accomplished using GATK’s SelectVariants with the peramaters: -window-size = 10, -QualByDept <2.0, -MapQual <40.0, -Qscore <100, -MapQualityRankSum <-12.5, -StrandOddsRatio > 3.0, -FisherStrandBias >60.0, -ReadPosRankSum <-8.0 for SNPS, and -window-size = 10, -QualByDepth <2.0, -MapQualityRankSum <-12.5, -StrandOddsRatio >4.0, -FisherStrandBias >200.0, -ReadPosRank <-20.0, -InbreedingCoeff <-0.8 for INDELS. Variants overlapping transposable elements were excluded from the variant pool by checking for overlap with the TE locations identified in the FungiDB v.46 release of the Af293 reference genome, using BEDtools -subtract (Quinlan & Hall, 2010). Variants were annotated with snpEff (Cingolani *et al*., 2012) based on the GFF annotation for Af293 v.46 from FungiDB.

### Population structure

Prior to analysis, the presence of clones in the dataset was assessed using the R package poppr v.2.9.0 (Kamvar *et al*., 2014) with the functions mlg() and clonecorrect(). No clones were detected. Initially, we assessed broad-scale population structure in *A. fumigatus* using the Bayesian clustering approach STRUCTURE implemented in fastStructure v1.0 (Raj *et al*., 2014) across 361,717 polymorphic sites (single nucleotide polymorphisms). VCF files were first converted into the plink format using PLINK v.1.90b3.38 (Purcell *et al*., 2007) before running fastSTRUCTURE using the simple prior with K values ranging from 1-15 over 30 independent iterations with specified seed values. For each independent iteration, marginal likelihood values of K were obtained by employing the ‘chooseK.py’ function in fastSTRUCTURE, and assessed for significant increases in marginal likelihood using ANOVA and the R package multcomp (Hothorn *et al*., 2008) with post hoc testing using Tukey tests and Bonferroni correction for multiple comparisons. To further assess population structure, we used Discriminate Analysis of Principle Components (DAPC) (Jombart *et al*., 2010) and subsequent clade mapping onto the phylogeny. DAPC was implemented in the R package adegnet v.2.1.3 (Jombart, 2008) on a random subset of 100K polymorphic SNP sites. The optimal number of groups (K) was identified based on Bayesian information criterion (BIC) score using the find.clusters() function in adegnet, evaluating a possible range of 1 to 15 clusters to identify the elbow of the BIC curve following (Jombart *et al*., 2010). To avoid overfitting, the optimal number of PCs retained in the DAPC was chosen using the optim.a.score() function, and determined to be PC=3 out of a possible PC range of 1-200. Evaluation of population sub-structure was carried out by iteratively running DAPC and fastSTRUCTURE on the three primary clades identified above. To do this, clade-specific VCF files were subset using VCFtools *-keep* (Danecek *et al*., 2011), and invariant sites were removed using the BCFtools *-view* command (Li, 2011), before running the pipeline as above. The fixation index (F_ST_) was calculated in the R package hierfstat v.0.5.9 (Goudet, 2005) using the dosage format to account for unequal population sizes (Weir & Goudet, 2017) and computed with the fst.dosage() function. Pairwise F_ST_ (the between population fixation index) was calculated in hierfstat using the fs.dosage()$Fst2×2 function. As a second measure of population differentiation, we ran analysis of molecular variance (AMOVA) using the R packages poppr and ade4 (Dray & Dufour, 2007) with the function poppr.amova() with method = “ade4”. Significant differentiation between populations was evaluated using a Monte-Carlo test with the function randtest() in poppr, over 999 iterations. To distinguish introgression from incomplete lineage sorting, we calculated Patterson’s D (ABBA-BABA) in Dsuite using the Dtrios tool (Malinsky *et al*., 2021), with significance determined using a 20 block-jackknife approach, with three populations and *A. fischeri* as the out group, on a VCF file prepared as above except with the addition of *A. fischeri*. Initially, the assembly for *A. fischeri* strain NRRL 181 was downloaded from FungiDB v.55. The assembly was then cut to simulate Illumina paired-end reads to a read depth approximately equal to the average for the *A. fumigatus* strains, and run as above.

### Linkage Disequilibrium

To estimate Linkage Disequilibrium (LD) Decay, the VCF SNP files (run without *A. fischeri*) generated above were used in conjunction with the clade assignments generated in DAPCA for K=3 to assess LD decay using PLINK (Purcell *et al*., 2007). LD decay was assessed both for all isolates together, and for all isolates separated by clade. Additionally, we assessed LD decay on a sample size rarified set of isolates at n=12 averaged over 20 independent runs. To do this, the samples were rarefied using a custom BASH script, where for each clade, random draws of 12 isolates per clade were taken over 20 iterations. Sampling did not allow for the same strain to appear more than once in a single iteration, but did allow sampling with replacement between iterations. To specifically evaluate the impact of sample size on LD estimates, we also ran the analysis for n = 5, 10, 20, 50 or 100 isolates (averaged over 20 independent runs) randomly drawn without regard to population structure. For all analysis, strains were subset (when appropriate) from the VCF file as noted above, using VCFtools *-keep* and BCFtools *-view* before converting VCF files to PLINK format using PLINK v1.90b3.38. Linkage Disequilibrium (LD) decay was estimated in PLINK using the squared genotypic correlation coefficient (r^2^) over all SNPs present in each group. We considered up to 99,999 variants per window and all r^2^ values to ensure fine scale resolution. Pairwise distance comparisons were limited to 500 kb (with parameters: --r2 –allow-extra-chr –ld-window-r2 0 –ld-window 99999 –ld-window-kb 500). Distance matrixes were constructed in BASH and mean r^2^ for each distance was assessed across each iteration, and then across each group to generate LD decay curves for each group using *ggplot2* (Whitmore, 2020). To estimate half-decay values (LD50) in BP for each dataset, we averaged the r^2^ for each position across all replicates in each group and calculated the r^2^ mid-point as (minimum r^2^ + (maximum r^2^ – minimum r^2^) / 2). To obtain LD50 in BP, we then calculated the x intersect of the r^2^ mid-point for each clade using the approx() function in R.

### Phylogenomics

Aligned SNPs from all isolates were used to construct a phylogeny employing the Maximum Likelihood algorithm IQ-TREE using the +ASC parameter to account for SNP based ascertainment bias (Nguyen *et al*., 2015). The best fit model according to Bayesian information criteria (BIC) score was assessed using the ModelFinder function in IQ-TREE and was determined to be GTR+F+ASC which was run over 1000 rapid bootstrap iterations. Individual branch support values were assessed using a Shimodaira-Hasegawa approximate likelihood ratio test over 1000 iterations. Tree rooting was determined by evaluation of the SNP tree run as above but including the outgroup *A. fischeri*. Based on outgroup position, the root was determined to be at the basal node separating Clades 2 and 3. Tree visualization and trait mapping was carried out using the R package *ggtree* (Yu, 2020).

### Distribution of mating type idiomorphs

To identify MAT type for each strain, we downloaded reference CDS DNA sequences for MAT1-1 (GenBank number AY898661.1 from strain AF250) and MAT1-2 (Afu3g06170, NCBI number NC_007196.1 from strain Af293) from NCBI. We constructed databases composed of all scaffolds for each genome, and ran BLASTN (Altschul, 1990) with the parameters -evalue .0001 and -word_size 10, against both MAT idiomorphs. BLAST hits were formatted as fastA files using a custom BASH script, and aligned using MAFFT v.7.471 (Katoh & Standley, 2013). To ensure that strains with significant alignments to both MAT1-1 and MAT1-2 were not artifacts in the *de novo* assembled scaffolds, these strains (n=11) were further evaluated by aligning the raw reads onto the MAT reference sequences. To accomplish this, reference sequences were indexed using Bowtie2 v2.3.4.1 (Langmead & Salzberg, 2012) with the -build function, and SAM files created with the -very-sensitive-local function. SAM files were converted to BAM using SAMtools -view function and sorted using the -sort function. Depth profiles of the alignments were created using the SAMtools -depth function and graphed in R using the package ggplot2. To create fastA files of the alignments, we used the BCFtools -mpileup function to create VCF files, indexed the VCF files using SAMtools -tabix, and used SAMtools -faidx to index the reference sequences. Then, using picard v2.18.3 we used the -CreateSequenceDictionary function to insert variant information for each strain into the reference MAT sequences, including “N”s for all positions where no alignments could be made. To assess the ploidy of these nine strains, we used whole genome K-mer analysis on forward reads implemented in the program Jellyfish (Marçais & Kingsford, 2011), and visualized using GenomeScope (Vurture *et al*., 2017), at K=21. To further assess the ploidy of these nine strains, we conducted allele frequency analysis on all heterozygous SNP sites aided by the R package vcfR (Knaus & Grünwald, 2017). Identification and mapping of the MAT1-2-4 gene (Afu3g06160) was conducted using BLASTP matches to Af293 gene assignments to the pan-genome clusters, and mapped onto the phylogeny as above.

### Presence-absence variance and SNP distribution of antifungal resistance genes and virulence factors

A database of characterized *A. fumigatus* antifungal resistance variants was assembled from the literature and combined with all MARDy database entries for *A. fumigatus* (http://mardy.dide.ic.ac.uk) (Table S2). Variant scans were conducted on all characterized resistance-related amino acid changes, available for six genes, as well as surveying all amino-acid changing variants across these six genes and an additional seven genes associated with azole resistance in the literature, but for which no regulatory amino acid changes have been characterized. Variant scans were conducted using custom scripts in R and mapped onto the phylogeny using the packages ggplot2 and ggtree. We used the BLASTP matches to Af293 gene assignments to validate the credibility of clade-specific gene family assignments and to look for presence – absence variation in *A. fumigatus* secondary metabolite biosynthetic gene clusters, consisting of 230 genes across 26 clusters as defined in (Bignell *et al*., 2016). To determine the boundary of a large deletion in the gliotoxin cluster covering *gliI*, *gliJ* and *gliZ*, in some strains, we extracted the genes up and down-stream from the *de novo* assemblies, Afu6g09690 and Afu6g09660 (*gliP*), using genome coordinates as defined for the Af293 reference genome. These sequences were then aligned onto Af293 using minimap (Li, 2016). To visualize the boundary of the deletion, a truncated example of *gliP* was extracted using samtools *faidx* (from AF100-12_9) using the coordinates identified in the alignment, and DNA was translated for both reference and truncated versions of the *gliP* gene using exonerate (Slater & Birney, 2005). Architecture was modeled using domain annotations for *gliP* from the Af293 reference in UniProt (UniProt Consortium, 2021), and visualized using the R package drawProteins (Brennan, 2018).

## RESULTS

### Genome sequencing and assembly statistics

Average sequencing depth per strain ranged from 12X coverage (IFM_59356-2) to 359X coverage (MO91298SB) with a mean depth of 75X over the 29 Mb *A. fumigatus* genome. BUSCO scores ranged from 96.3% (MO91298SB) to 99.4% (16 strains) completeness with a mean of 98.9%. The total number of scaffolds ranged from 38 (Afum_84-NIH) to 872 (AF100-1_3) with a mean of 191. L50 ranged from 3 to 4, with a mean of 3.9. N50 ranged from 3,530,065 bp to 4,348,233 bp with a mean of 3,935,233 bp. Genome size ranged from 27,437,509 bp (CM2495) to 31,568,453 bp (NCPF-7820) with a mean of 29,127,620 bp (Table S1).

### Population Structure

fastSTRUCTURE’s marginal likelihood values increased until K= 5, but did not increase significantly after K=4 with a mean marginal likelihood difference of only 0.0005 between K=3 and K=4 (Fig. S1). Because fastSTRUCTURE tends to overestimate K when K is small (Raj *et al*., 2014), and is predicated on the assumption of locus independence due to recombination, which is violated by clonality, we further defined K using DAPC analysis. The optimal K was approximately 3 (Fig. 1A), using the BIC criteria with three principal components retained (Fig. 1B). DAPC supported clear separation of three primary clades (Fig. 1C). Clade 1 was the largest with 200 strains, followed by Clade 2 with 45 strains and Clade 3 with 15 (Table S3). According to DAPC, none of the strains in this study met the criteria for admixture (membership coefficients < 0.85), as membership coefficients were essentially 1 in all cases (Table S3). While clade assignment to one of the three clades using fastSTRUCTURE placed all strains in the same respective clades as DAPC, fastSTRUCTURE membership coefficients were lower overall than DAPC membership coefficients (Fig. 1D, Table S4) and a total of 8 strains had membership coefficients < 0.85, indicative of admixture or incomplete lineage sorting between 2 or more of the clades (Fig. 1E). These included three strains (RSF2S8, 12-7505220, and 10-01-02-27) assigned to Clade 2 but with ancestry from Clade 1, and five strains (F7763, CF098, F18304, CM7632 and the reference strain AF293) assigned to Clade 1, but with ancestry from both Clades 2 and 3. To investigate population substructure within the three clades, we used iterative DAPC and fastSTRUCTURE analysis of on each clade separately. Subset VCF files excluding all invariant sites consisted of a total of 61,618 SNPs in Clade 1, 132,130 SNPs in Clade 2 and 155,585 SNPs in Clade 3. The difference in SNP abundance between the three clades are consistent with the location of the reference Af293 strain in Clade 1. The optimal number of PCs for the sub-clades was determined to be 5 for the substructure within Clade 1 (Fig. S2A), and 3 for the substructure within Clade 2 (Fig. S2B). The optimal number of PCs for Clade 3 was determined to be 1 (Fig. S2C), and was therefore not further analyzed with DAPC. Iterative DAPC analysis supported 5 sub-clades within Clade 1, and 5 sub-clades within Clade 2 (Fig. S2D-E, Table S5). DAPC and fastSTRUCTURE returned slightly different clade memberships for sub-clades at K=5 (Table S5-S6). fastSTRUCTURE analysis at K=5 supported frequent introgression between sub-clades in both Clade 1 (87 out of 200 strains) (Fig. S2F), and Clade 2 (7 out of 45 strains) (Fig. S2G).

**Fig. 1:**
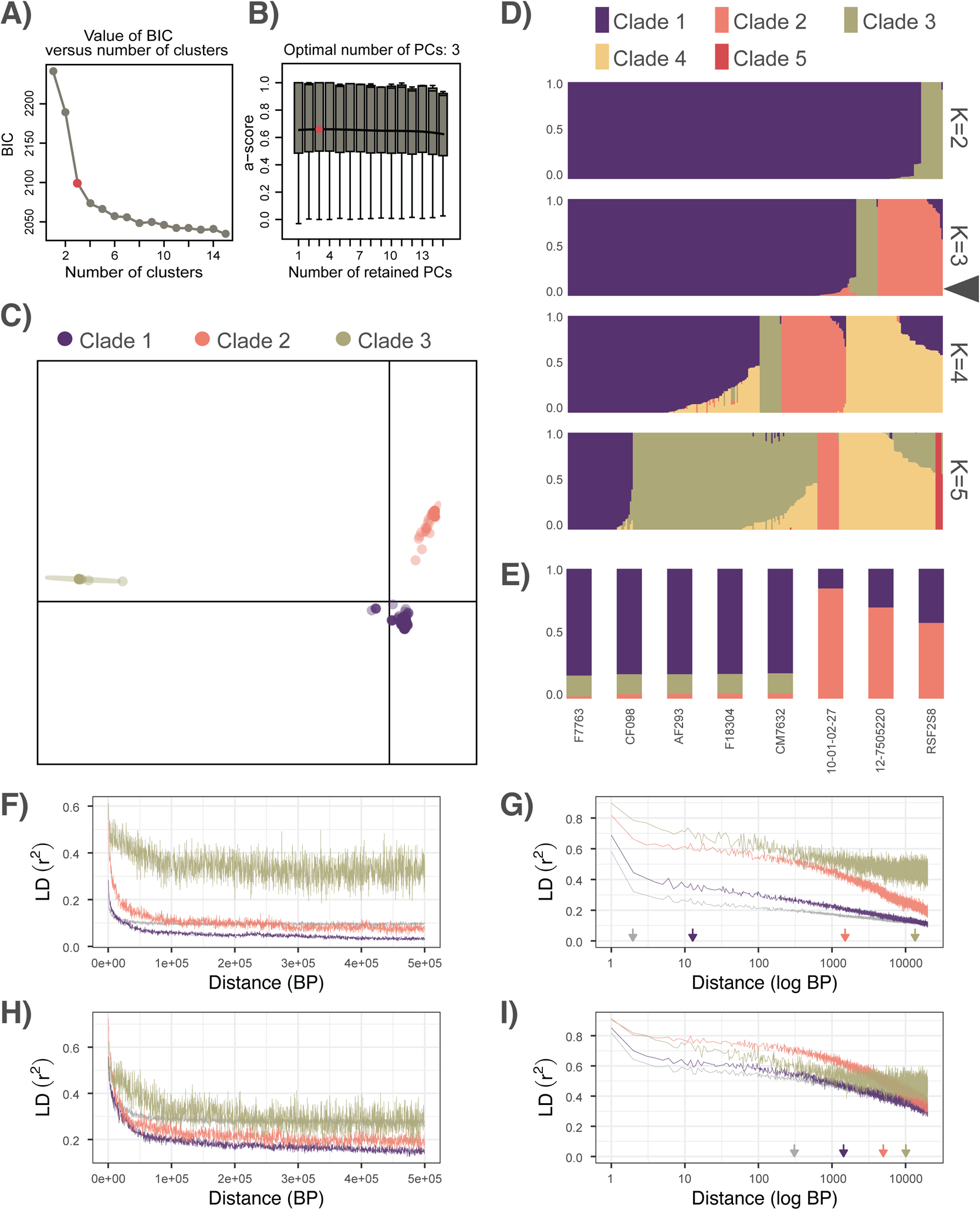
*A. fumigatus* population structure. **A)** Bayesian Information Criterion (BIC) score vs. number of clusters, with elbow of the curve at K=3 clusters **B)** choice of PCs for DAPCA was found to be optimal at PC=3 according to a-score **C)** DAPCA plot depicting three primary populations with clear separation between clusters. The x axes represents the first discriminant function, y axes represents the second discriminant function, ellipses represent 95% confidence areas. **D)** STRUCTURE plots of K=2 to K=5 populations support low levels of admixture between clades at K=3 (highlighted with triangle). **E)** Isolates with < 0.85 probability of assignment to a single clade, indicative of admixture or incomplete lineage sorting between clades. **F-G)** Linkage disequilibrium decay (r^2^) for all isolates in each respective clade, or all isolates across all clades (in grey) **F)** Linear-linear plot of LD decay. **G)** Zoomed-in log-linear plot of LD decay, with arrows indicating LD50 (half decay) values. **H-I)** Linkage disequilibrium decay (r^2^) for sample size normalized isolates (n=12 isolates, averaged over 20 draws) in each respective clade, or for all isolates across all clades (in grey) **H)** Linear-linear plot of LD decay. **I)** Zoomed-in log-linear plot of LD decay, with arrows indicating LD50 values.

The value of Patterson’s D was 0.135 (Z = 24.32, *p* < 0.0001, Dmin = 0.118), indicating introgression over incomplete lineage sorting. Mapping the three clades onto the phylogeny was congruent with clade membership, with all clades demonstrating monophyly, with the exception of all of the Clade 1 strains with signatures of admixture, which clustered on their own branch between Clades 2 and 3. Overall F_ST_ was 0.824, and within population F_ST_ was 0.767, 0.825, ∼0.880 for populations 1, 2 and 3 respectively. Pairwise (between population) F_ST_ was 0.531 between Clade 1 and 2, 0.887 between Clade 2 and 3, and 0.859 between Clade 1 and 3. Similarly, AMOVA supported that the majority of variation, 69.8%, occurred between populations (σ = 45,009.85, p = 0.001) whereas differences between individuals in the populations accounted for only 30.2% of the total variation (σ = 19,482.91). LD50 estimates were extremely low when evaluated without considering clade or sample size (1.97 BP considering all 260 isolates), and notably higher when taking clade into account (12.79 BP for Clade 1, 1,508.32 BP for Clade 2 and 13,474.24 BP for Clade 3, Fig. 1 F-G). The influence of sample size on LD estimates showed a strong inverse relationship between sample size and LD (Fig. S3). Similarly, sample size normalized LD estimates at n=12 were notably higher overall (LD50 = 308.00 BP) and for Clades 1 (LD50 = 1,425.24 BP) and 2 (LD50 = 4,974.58 BP), but not for the smallest and most closely related clade, Clade 3 (LD50 = 10,096.73 BP) (Fig. 1 H-I).

### Pan-genome

OrthoFinder identified 15,309 gene families including 8,866 core genes (57.91% of the total), 4,334 accessory genes (28.31%), and 2,109 singletons (13.78%). PIRATE identified 15,476 gene families, including 8,600 core genes (55.57% of the total), 3,618 accessory genes (13.92%), and 3,258 singletons (21.05%) (Fig. S4A). The distribution of the number of genomes represented in each gene family was characteristically U-shaped, and ranged between 1 and 260 for both OrthoFinder (Fig. 2B), and PIRATE identified gene families (Fig. S4B). Gene accumulation curves suggested a semi-open genome structure where accessory gene families excluding singletons leveled after sampling ∼150 genomes, but continued to increase when considering singletons along with accessory genes for both OrthoFinder (Fig. 2C) and PIRATE identified gene families (Fig. S4C). The number of unique accessory gene families per strain from OrthoFinder identified gene families ranged from 692 in AF100-12_21, to 1,450 in IFM_60237 with an average of 1163 per strain (Fig. 2D). PIRATE identified gene families ranged from 665 in AF100-12_21 to 1,376 in IFM_60237 with an average of 974 per strain (Fig. S4D). The number of unique singleton gene families per strain ranged from 1 (34 strains) to 130 in the strain AFUG_031815_1869, with an average of 8 per strain. One strain (AF100_12_23) contained no singleton gene families (Fig. 2E). PIRATE identified gene families supported singleton gene families ranging from 1 (27 strains) to 191 (in AFUG_031815_1869) with an average of 13 per strain, and 45 strains with no singleton gene families (Fig. S4E). Although AFUG_031815_1869 was an outlier in the number of singletons present in both the OrthoFinder and PIRATE analysis. All AFUG_031815_1869 genome quality metrics were average relative to the rest of the dataset (200 scaffolds, average depth of 53bp, and a BUCO score of 99.3), and it did not appear that accessory genes were being miscategorized as singletons, so we elected to include this strain in all downstream analysis. Overall, both accessory (*p* = 8.3e^-8^, R^2^ = 0.12) and singleton *p* = 4.9e^-5^, R^2^= 0.06) gene families were significantly related to genome size, although with low R^2^ values, particularly for singleton gene families (Fig. S5). Neither the number of accessory gene families per strain, nor the number of singleton gene families per strain was significantly different between clades, with means of 1,171, 1,160, and 1,075 accessory gene families per genome in Clades 1, 2 and 3 respectively, and 9, 4, and 8 singleton gene families per genome for OrthoFinder identified gene families. These results were largely consistent for PIRATE identified gene families, with means of 980, 973, and 909 accessory gene families per genome in Clades 1, 2 and 3 respectively, and 14, 6, and 16 singleton gene families per genome, and no significant differences in gene family abundance between clades. Because of the relatively similar results obtained between OrthoFinder and PIRATE, we conducted the remainder of the analysis solely on the OrthoFinder identified gene families (hereafter referred to as just gene families). Clade 1 contained 1,256 unique accessory gene families (not found in the other two clades) (average 6 per strain), Clade 2 contained 95 unique accessory gene families (average 2 per strain), and Clade 3 contained 115 unique accessory gene families (average 8 per strain) (Fig. 3A). Clade defining gene family gains (defined as present in >90% of strains in the clade, but in 0 strains from the other two clades) were 0 for Clade 1, 2 for Clade 2 and 23 for Clade 3. Analysis of clade-specific gene family absences identified 24 gene families missing in Clade 1 (with 5 Clade-defining absences, defined as lost in all isolates of Clade 1 but present > 90% of all isolates from Clades 2 or 3) (Fig. 3B), as well as 270 gene families absent in Clade 2 (with 25 clade-defining absences), and 991 gene families absent in Clade 3 (with 125 Clade-defining absences).

**Fig. 2:**
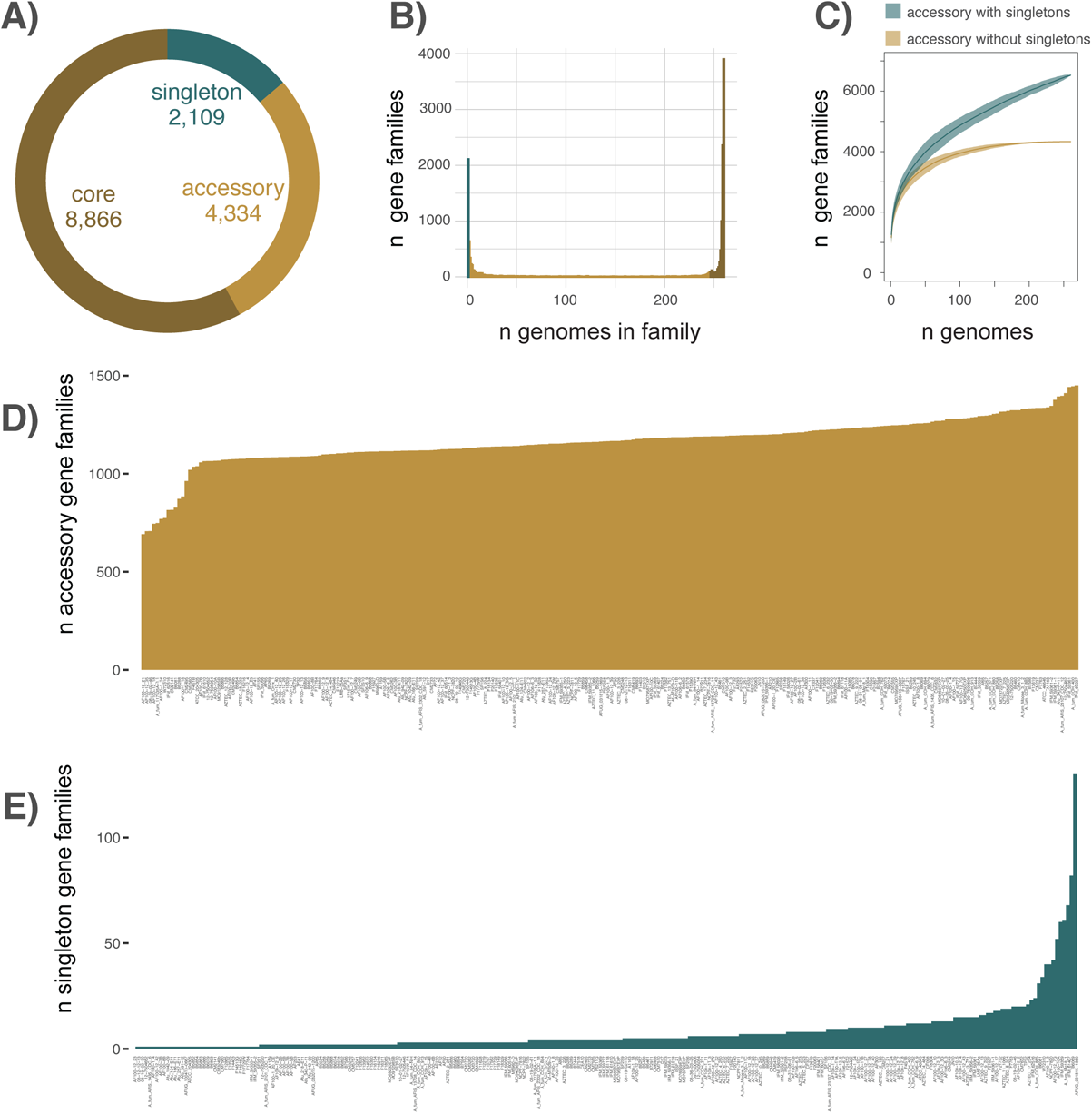
Pan-genome gene family distribution. **A)** The pan-genome of 260 *A. fumigatus* strains included 15,508 gene families in total, including 8,595 (55.42%) core genes (present in >95% of strains), 3,660 (23.60%) accessory genes (present in >2 and < 248 strains) and 3,253 (20.98%) singletons (present in only one isolate). **B)** The distribution of the number of genomes represented in each gene family. **C)** Gene family accumulation curves, including (green) and excluding (yellow) singletons. **D)** The distribution of unique accessory gene families by strain. **E)** The distribution of unique singleton gene families by strain.

**Fig. 3:**
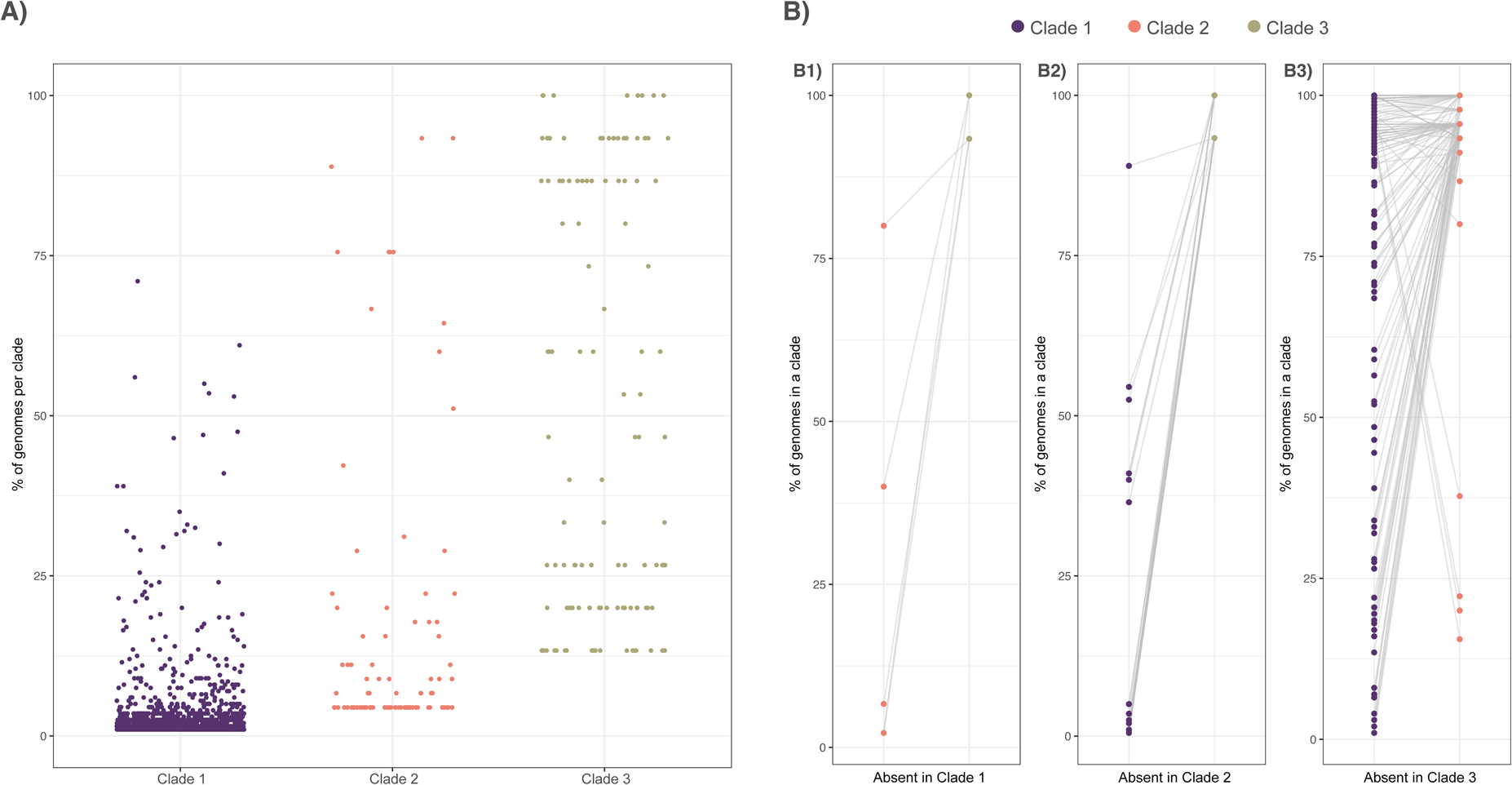
Distribution of clade-specific gene family gains and absences. **A)** The occurrence of Clade-specific gene families in that Clade (x-axis), defined as genes that exist in that Clade in more than one genome (singletons were excluded), by prevalence in that Clade (y-axes), and highlights that while many Clade-specific gene families that exist in a low number of isolates, others are present in nearly all the isolates of that clade and in no other isolates. The distribution of occurrence frequency varies between the three Clades, due to differences in Clade size, with 1,256 accessory gene families exclusive to Clade 1, 95 exclusive to Clade 2 and 115 exclusive to Clade 3 **B)** The occurrence of Clade-specific gene family absences for clade defining absences, where each panel depicts the occurrence of each gene in the two clades where the gene is not absent. The distribution of each gene is connected by a grey line **B1)** Gene families lost in all isolates of Clade 1 but present in greater than 90% of the isolates in either Clade 2 or Clade 3 (n = 5) **B2)** Gene families lost in all isolates of Clade 2 but present in greater than 90% of the isolates in either Clade 1 or Clade 3 (n = 25) **B3)** Gene families lost all isolates of Clade 3 but present in greater than 90% of the isolates in either Clade 1 or Clade 2 (n = 125).

The proportion of secreted proteins was significantly higher in the core gene families than for accessory or singleton gene families (pairwise proportion test at *p* < 0.05). No significant differences were observed in the proportion of secreted proteins between the three clades (Fig. S6). Significant BLASTP hits for representative gene family sequences were found for 9,512 of the 9,840 genes currently annotated in the Af293 reference genome Because the Af293 reference strain is a member of Clade 1, Clade specific gene families in Clades 2 and 3 should not be present in our BLAST searches if they are truly clade specific. Clade specific gene families present in Clade 2 had no significant BLAST hits to the reference strain, but Clade 3 specific gene families had two gene families with significant matches to genes in the Af293 reference (Afu8g01650 and Afu3g02860 – both encoding proteins of unknown function), indicating low levels of sequence similarity among genes that our approach classified as unique families. Clade-specific gene families in Clade 1 had 28 gene families that could be assigned an Af293 reference gene ID, all other Af293 annotated gene families were present in more than one clade (Table S7).

### Distribution of mating type idiomorphs

Whereas the MAT1-1 idiomorph has a unique sequence structure encoding the α-box domain, MAT1-2 contains both a unique region encoding a high-mobility group (HMG) region and a region conserved between mating types (Paoletti *et al*., 2005). Additionally, MAT1-2 strains are expected to contain the gene MAT1-2-4 (Afu3g06160), essential for sexual recombination (Yu *et al*., 2017). We therefore expected MAT1-1 strains to have full-length alignments to the MAT1-1 reference and a truncated alignment to the MAT1-2 reference, and MAT1-2 strains to have full-length alignments to the MAT1-2 reference, no alignment to the MAT1-1 reference, and the presence of MAT1-2-4. Using this criteria, we identified 144 isolates in our set with the MAT1-1 idiomorph and 105 isolates with the MAT1-2 idiomorph (Fig. 4), a ratio of 48:35. Additionally, 11 isolates had significant alignments to both MAT1-1 and MAT1-2, hereafter referred to as ‘unknown’ mating type. All unknown mating types also contained the MAT1-2-4 gene. All three clades contained both MAT types, in MAT1-1:MAT1-2 ratios of 107:85 (8 unknown) in Clade 1, 26:16 (3 unknown) in Clade 2 and 11:4 (0 unknown) in Clade 3. The 11 isolates with unknown mating type were further examined using read mapping and read depth analysis, which confirmed that for each of the nine strains, raw reads did indeed map onto the full length of both reference idiomorphs, however, alignment depths of MAT1-1 relative to MAT1-2 differed among strains, and only three strains displayed relatively equal depth profiles for both MAT-1 and MAT-2 (AF100-1_3, IFM_59359, and IFM_61407) (Fig. S7). To investigate the possibility that these 11 strains represented diploids, we first conducted whole genome K-mer analysis, and found that nine of the 11 strains had strong single peaks supporting haploidy, while two strains, AF100-1_3 and IFM_59359, had moderate secondary peaks, potentially indicative of diploidy (Fig. S8). To further assess strain ploidy, we conducted allele frequency analysis on all heterozygous SNPs, and found that three strains (AF100-1_3, IFM_59359, and IFM_61407) displayed notable peaks at 1/2 frequency, lending further support to the potential for diploidy (Fig. S9). Three strains (08_36_03_25, NCPF_7816, and SF2S9) had no significant peaks at ½ allele frequency, but did display moderate shoulders sloping down from 1 (representing SNPs with base calls different than the Af293 reference) and up to 0 (representing SNPs with the same base call as the reference) potentially indicative of noise in the sequencing (miscalls, contamination or tag switching), or low levels of copy number variation or segmental duplication. The remaining five strains (AF100-1_18, AF100_12_5, Afu_343_P_11, B7586_CDC_30, and AF100_12_7G) displayed only low levels of 1/2 allele frequency but did display paired peaks below 1/4 and above 3/4, indicative of haploidy with large scale copy number variation or segmental duplication in these strains (Knaus & Grünwald, 2018).

**Fig. 4:**
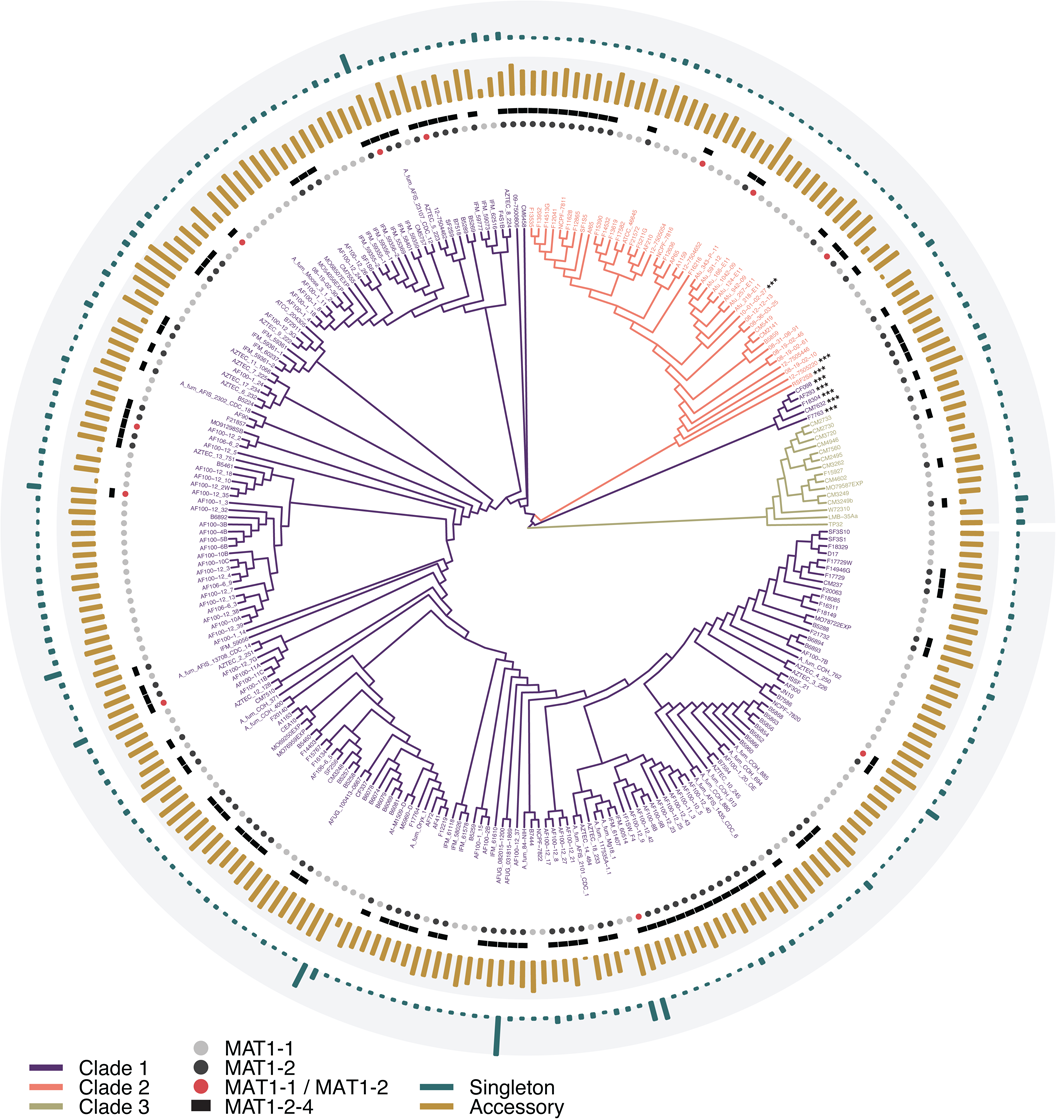
Phylogeny of *A. fumigatus* mapped against pan-genome distribution and MAT type. Overall abundance of accessory and singleton gene families were not structured by clade. The distribution of MAT type was random across the phylogeny, with both mating types present in all three clades. MAT idiomorphs were not present in equal proportion, with 144 strains containing the MAT1-1 idiomorph, 105 containing MAT1-2, and 11 strains presenting significant alignments to both MAT1-1 and MAT1-2. All strains with the MAT1-2 idiomorph also contained the MAT1-2-4 gene, including the 11 strains with alignments to both MAT1-1 and MAT 1-2. Strain names with the *** suffix denote the 8 strains demonstrating significant admixture between two or more clades.

### Functional analysis

While only 1.75% of core gene families were unable to be assigned with any functional annotation, this number was greater for accessory (20.52%) and singleton (18.73%) gene families. The percent of gene families unable to be assigned functional annotation was similar between Clades 1 and 2 (18.47% and 18.95% respectively) but higher for Clade 3 (39.13%). Gene Ontology enrichment analysis of gene families found exclusively in the core, accessory and singleton categories identified significant GO terms in all three categories. The same analysis targeting clade-specific gene accessory gene families, identified terms significantly enriched in Clades 1 and 3 (Fig. 5). Clade 2 was not significantly enriched for any GO terms. core gene families were generally enriched for terms related to housekeeping functions like transport, signal transduction and general cellular processes (Fig. 5A). Accessory gene families were enriched in terms for nitrogen, carbohydrate and phosphorus metabolic processes, with a small number of terms associated with molybdoprotein metabolic processes (Fig. 5B). Singleton gene families were enriched for terms relating to carbohydrate and nitrogen metabolism, as well as primary metabolism and transcriptional regulation (Fig. 5C). The most significantly enriched terms in Clade 1 also included terms associated with metabolism, including carbohydrate and nitrogen processing, as well as gene families associated with transmembrane transport and vesicle mediated transport (Fig. 5D). There was only one term significantly enriched in Clade 3, also for carbohydrate metabolism (Fig. 5E).

**Fig. 5:**
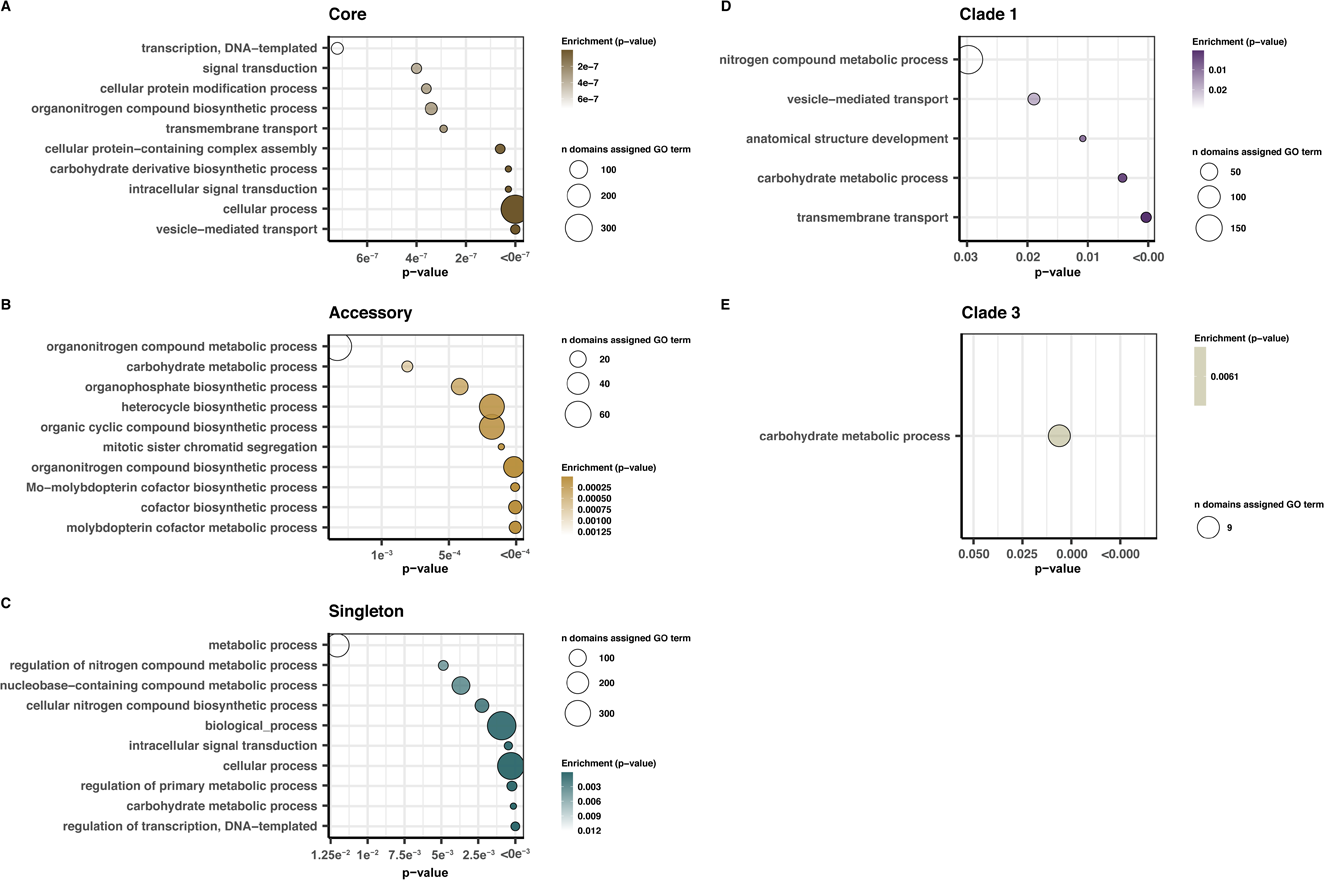
GO enrichment. Significantly enriched Biosynthetic Process Gene Ontology (GO) terms associated with accessory gene families present exclusively in different *A .fumigatus* pan-genome categories **(A-C)** and clades **(D-E)**. Enrichment analysis was conducted for each experimental set against a background of all GO terms for all gene families in the *A. fumigatus* pan-genome. p-values represent results of Fisher-exact test of the top ten most significantly enriched terms (for terms enriched at *p* < 0.05) **A)** core, **B)** accessory, **C)** singleton, **D)** Clade 1, **E)** Clade 3. Clade 2 had no significantly enriched terms and is not shown here.

To further investigate metabolic genes differentially abundant between the three clades, we used gene families with Af293 reference annotations that fell into GO categories for nitrogen, carbohydrate and phosphorus metabolism. We identified 25 genes with Af293 GO annotations for the term organonitrogen compound metabolic process that were significantly differently abundant between the three clades (Fig. S10A). Additionally, there were seven genes with differential abundance for the term carbohydrate metabolic process (Fig. S10B), and five genes differentially abundant for the term organophosphate biosynthetic process (Fig. S10C).

### Distribution of CAZymes

In total, 139 different CAZyme classes were identified across the *A. fumigatus* pan-genome. The CAZyme counts per genome averaged 460 (sd=7.7, min=422 in F21732, max=478 in IFM_59361) (Table S8). Twenty-eight of these CAZyme classes displayed patterns of differential abundance between the clades (Fig. 6A, Table S9). These abundance profiles represented both copy number variation and clade-specific gene gains and absences in families of Auxiliary Activities (AAs, n=4), Carbohydrate-Binding Modules (CBMs, n=3), Carbohydrate Esterases (CEs, n=3), Glycoside Hydrolases (GHs, n=13), Glycosyl Transferases (GTs, n=4).

**Fig. 6:**
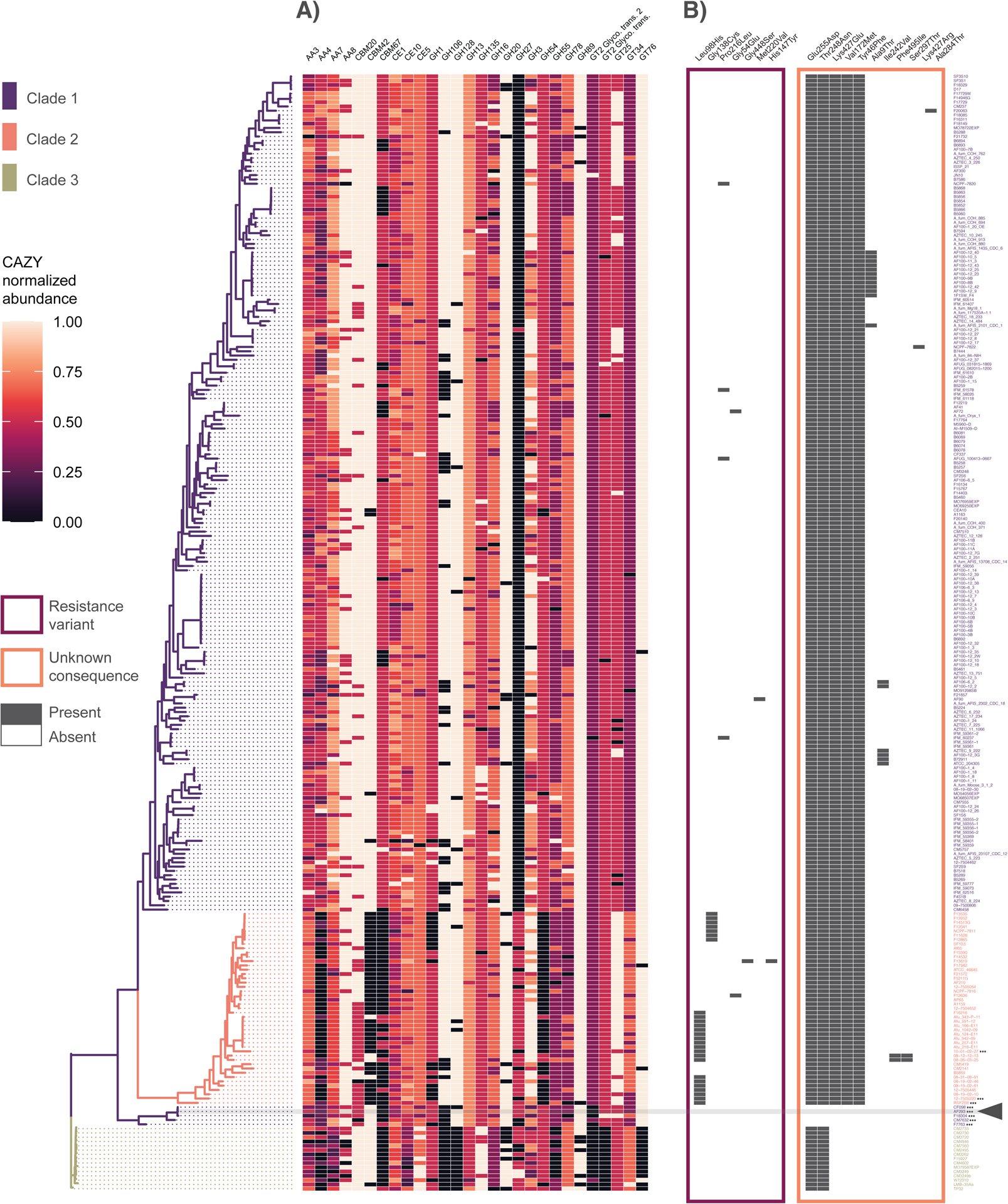
Distribution of CAZymes and Occurrence of *cyp51A* mutations across the *A. fumigatus* phylogeny. **A)** CAZyme profiles display patterns of both Clade-specific gene gains and Clade-specific gene absences across 28 CAZyme classes. CAZyme classes with differential abundance between the three Clades were determined by Kruskal Wallace tests at *p* < 0.001 after Bonferroni adjustment for multiple comparisons. Gene counts are normalized on a 0-1 scale (by CAZyme class) for visualization. **B)** Identification of non-synonymous variants in the *cyp51A* gene across the phylogeny demonstrated structured occurrence of both known resistance variants (framed in pink), and variants with unknown functional impacts (framed in orange). Reference strain Af293 is highlighted in grey and with triangle. While the Leu98His variants (corresponding to the azole resistant TR34/Leu98His genotype) as well as azole resistant variant Gly138Cys, were found exclusively in Clade 2, other characterized variants were scattered in low abundance throughout Clade 1, 2 or both. No characterized resistance variants were found in Clade 3. While some non-synonymous variants with unknown functional impacts occurred frequently in *cyp51A*, and represented changes in a single branch leading to the reference strain Af293 (Glu255Asp, Thr248Asn), others were absent form this branch and absent in Clade 3 (Lys427Glu, Val172Met, Tyr46Phe), while others were low abundance and exclusive to Clade 1 (Ala9Thr, Lys427Arg, Ile242Val, Ala284Thr), or Clade 2 (Ser297Thr, Phe495Ile).

### Presence-absence variance and SNP distribution of antifungal resistance genes and virulence factors

Variant scans of 27 characterized amino acid changing mutations across six genes associated with resistance identified variants only in the *cyp51A* gene. In *cyp51A*, a total of seven resistance variants were identified occurring in a total of 34 strains (Table 1, Fig. 6). The most abundant of these was the TR34/Leu98His genotype (no strains with TR46/Tyr121Phe/Thr289Ala genotype were found). All strains with the TR34/Leu98His genotype, as well as all strains containing the azole resistant variant encoding Gly138Cys, were found exclusively in Clade 2. Other low abundance variants representing characterized *cyp51A*-mediated azole resistance were scattered throughout Clade 1 (Pro216Leu, Met220Val) or Clade 2 (Gly448Ser, His147Tyr), or represented in both Clade 1 and Clade 2 (Gly54Glu). No characterized *cyp51A* resistance variants were found in Clade 3. An additional scan for all non-synonymous *cyp51A* variants revealed 11 additional sites with unknown functional impacts. Two of these sites occurred frequently across the phylogeny, and represented changes in a single branch containing Af293 and the other introgressed isolates in Clade 1 (Glu255Asp, Thr248Asn), while three were absent from both the Af293 branch and all Clade 3 isolates (Lys427Glu, Val172Met, Tyr46Phe). Six additional sites occurred in low abundance and were either exclusive to Clade 1 (Ala9Thr, Lys427Arg, Ile242Val, Ala284Thr), or Clade 2 (Ser297Thr, Phe495Ile). Other amino acid changing variants in genes associated with resistance, but where the functional consequence of these variants has not been characterized, were found in all other genes investigated, including AFUA7G01960, *artF*, *cdr1B/abcC*, *<u>cox10</u>*, *cyp51B*, *fks1*, *hapE*, *hmg1*, *mdr1, mdr2, mdr3*, and *mdr4* (Fig. S11). Similarly, these variants of unknown consequence at times represented the dominant allelic state (where the Af293 reference is not representative of the population) and were at other times structured by clade (such as the AFUA7G01960 Leu76Phe variant occurring almost exclusively in Clade 2, multiple the clade-specific variants in *mdr4* in Clade 3 isolates).

**Table 1:** Occurrence summary of *cyp51A* variants by clade. Seven non-synonymous variants associated with antifungal drug resistance were identified in the *cyp51A* gene, representing 34 isolates. Percentages represent the total number of a given amino acid change across all three clades out of 261 isolates scanned.

|  | Leu98His | Gly138Cys | Pro216Leu | Gly54Glu | Gly448Ser | Met220Val | His147Tyr |
| --- | --- | --- | --- | --- | --- | --- | --- |
| Clade 1 | 0 | 0 | 4 | 1 | 0 | 1 | 0 |
| Clade 2 | 19 | 7 | 0 | 1 | 1 | 0 | 1 |
| Clade 3 | 0 | 0 | 0 | 0 | 0 | 0 | 0 |
| Total | 19(7.25%) | 7(2.67%) | 4(1.52%) | 2(0.76%) | 1(0.38%) | 1(0.38%) | 1(0.38%) |

We further used the Af293 gene annotations assigned to gene families to investigate gene presence-absence variation across 26 notable *A. fumigatus* secondary metabolite biosynthetic gene clusters (BGC, as defined in (Bignell *et al*., 2016)) (Fig. S12). Out of the 230 genes investigated, seven were convoluted with one other gene in the same cluster, where both genes clustered into the same gene family. These included Afu3g13700/Afu3g13690 in BGC 10, Afu4g00220/Afu4g00210 in BGC 13, Afu4g14560/Afu4g14550 in BGC 14, Afu6g09730/Afu6g09720 in BGC 20, Afu6g13980/Afu6g13970 in BGC 22, Afu8g00490 Afu8g00480 in BGC 25, and Afu8g02380/Afu8g02360 in BGC 26. Three genes (Afu1g10275 in BGC 2, Afu7g00140 in BGC 23 and Afu8g00450 in BGC 25) could not be confidently assigned to a gene family. We found substantial variation in conservation across the 26 clusters, with the lowest conservation in BGC 1 (Uncharacterized polyketide – with 259 strains missing at least one gene), BGC 12 (Uncharacterized non−ribosomal peptide, 240 strains missing at least one gene), BGC 16 (Uncharacterized non−ribosomal peptide−like, 134 strains missing at least one gene) and BGC 13 (Endocrocin, 112 strains missing at least one gene). Additionally, Afu8g00400 in cluster 25 could only confidently be called in one isolate (AF100-1_24). Eleven BGCs demonstrated high levels of conservation, with less than 10 strains missing any gene in the cluster. The most conserved was BGC 24 (Fumitremorgin) for which all genes were present in all strains, followed by BGC 8 (Fusarine C) with genes missing in only 3 strains, BGC 11 (Uncharacterized polyketide) with genes missing in only 4 strains, BGCs 21 (Fumiquinozalines) and 3 (Ferricrocin), with genes missing in only 5 strains, BGC 7 (Uncharacterized polyketide) missing genes in only 6 strains, BGCs 6 (Fumigaclavine), 9 (Hexadehydroastechrome), and 19 (Uncharacterized non−ribosomal peptide), all missing in only 7 strains, BGC 5 (DHN Melanin) missing in only 8 strains, and BGC 23 (Neosartoricin) missing in 9 strains. We further investigated the likely implications of gene loss in the 13 genes encoding the gliotoxin cluster (BGC 20) across the *A. fumigatus* phylogeny. These 13 genes were assigned to 12 gene families (with *gliF* and *gliN* both assigned to the same family). All 12 were highly conserved with low levels of gene absences. An exception to this finding was the wholesale loss of *gliI*, *gliJ* and *gliZ* from 8 strains clustered in Clade 1 (Fig. S12). All strains represent isolates from a long-term infection of a Cystic Fibrosis patient (Ross *et al*., 2021). Because *gliI*, *gliJ* and *gliZ* are immediately adjacent in the Af293 reference genome, we further investigated the boundaries of this deletion, and found that it encompasses a total 18,261 base pairs (bp), spanning Chr6 2336977..2355238 in Af293 reference coordinates. This deletion included 454bp of the gene Afu6g095906, whole gene losses of the genes Afu6g09600, Afu6g0910, Afu6g09620, Afu6g09630 (*gliZ*), Afu6g09640 (*gliI*), Afu6g09650 (*gliJ*) and 2619bp of Afu6g09660 (*gliP*) (Fig. S12B).

## DISCUSSION

Contrary to models of panmixia, we found strong evidence for three distinct populations of *A. fumigatus*, with secondary population structure in both Clade 1 and Clade 2. This structure was supported by both DAPCA clustering and F_ST_ values both overall and between populations, as well as by AMOVA analysis (Fig. 1 A-C). Differentiation was lower between populations 1 and 2, with population 3 more divergent. To assess recombination ancestry between the clades, we first performed a STRUCTURE analysis (Fig.1 D), and identified only eight strains with clade assignment values indicative of mixed ancestry between two or three of the primary clades (Fig. 1 E). These included three strains assigned to Clade 2 and five strains assigned to Clade 1, including the Af293 reference strain. All five strains with mixed ancestry assigned to Clade 1 clustered together on the same branch which was positioned apart from the rest of Clade 1 and situated between Clades 2 and 3, typical of mixed ancestry (Fig. 4). Similarly, Patterson’s D supported a history of occasional introgression between clades over incomplete lineage sorting. Iterative DAPC and fastSTRUCTRE analysis identified approximately five sub-clades in both Clades 1 and 2 (Fig. S2D-E). Interestingly, while evidence for introgression between the three primary clades was limited to a few strains, there appeared to be substantial within-clade introgression between the sub-clades (Fig. S2F-G). We assessed recombination frequency using LD decay (a pairwise site comparison of the decrease in LD across genome space) and found that LD50 was extremely low (1.97 BP) when assessed across all isolates. Because LD estimates can be biased by ignoring population structure or unequal sample size, we also conducted LD assessments for each clade individually and over-all for different values of n-samples. We found that LD varied for each clade (lowest for Clade 1 and highest in Clade 3), both when considering all isolates in each clade and when normalizing by sample size. Sample-size normalized numbers generally increase the overall LD estimates (from 1.97 BP to 308.08 BP across all isolates). Given the sensitivity of LD to sample size and genetic structure, the normalized results likely offer a more acurate relative estimate of recombination frequency between the three clades. Overall, the range of LD values identified here demonstrate exceptionally high levels of recombination in *A. fumigatus*. For comparison, *Candida albicans* (considered to be obligately asexual) has reported LD50 of 162,100 (Nieuwenhuis & James, 2016). Conversely, *A. flavus*, which demonstrates relatively frequent outcrossing in some populations, has LD50 values estimated between 1,000-12,300 (Drott *et al*., 2020). Recent estimates of recombination rate have singled out *A. fumigatus* as potentially having a higher number of crossovers per meiotic event than any Eukaryotic species investigated to date (Auxier *et al*., 2022). Extremely high recombination rates would help to explain the extremely low LD-decay values, and exceptionally large pan genome size observed here.

Due to the high dispersibility and assumed substrate generalism in *A. fumigatus*, the species was originally thought to represent a single homogenous population (Pringle *et al*., 2005; Rydholm *et al*., 2006). However, recent investigations into *A. fumigatus* population structure have found mixed evidence for population stratification; a study using 255 *A. fumigatus* isolates from the Netherlands, and 20 genome-wide neutral markers, found evidence for five populations, with significant sexual recombination occurring in only a single population. Two studies employing 9 microsatellite markers and a large number of isolates (2,026 and 4,049 respectively), found mixed evidence for population stratification. The first spanned 13 countries, identified eight populations, and found evidence for both recombination and clonal reproduction, as well as evidence of gene flow between both domestic and intercontinental populations (Ashu *et al*., 2017). The second study included isolates from 26 countries and found evidence for only two primary populations, with possible underlying population substructure (Sewell *et al*., 2019). A whole genome sequencing approach using 24 isolates from three countries, defined *A. fumigatus* populations based on country of isolation, and concluded that *A. fumigatus* is panmictic, with as much total genetic diversity between domestic isolates, as between isolates form different continents (Abdolrasouli *et al*., 2015). Importantly, tests for recombination that do not take population structure into account, or incorrectly infer the number of underlying populations, can bias estimates of recombination rate (Yu *et al*., 2006).

By constructing *de novo* assemblies, we identified 5,180 more gene families than there are genes currently predicted in the Af293 reference genome (15,309 identified here vs. 9,840 protein coding genes, and 289 non-coding RNA genes currently predicted in Af293 in FungiDB as of October 2021). Because gene families represent clusters of homologues, including both orthologues and paralogues, genes and gene families are not expected to corelate in a one-to-one ratio, and the number of true genes in the *A. fumigatus* pan-genome is likely even higher than the number of gene families reported here. Method validation for our pipeline of *de novo* assembly and pan-genomic clustering, identified 10,167 genes (in 10,005 gene families) in the re-sequenced reference strain AF293, 38 more than are currently predicted in the curated Af293 reference strain. The small discrepancy between the number of genes identified in the pan-genomic analysis over those currently annotated in the reference highlights the high fidelity of the methods used here, where the additional genes identified for AF293 may represent either low levels of error in gene calling, or genuine variation between the reference Af293 isolate and the re-sequenced “AF293” isolate (Colabardini *et al*., 2022).

The majority of gene families were part of the core pan-genome of *A. fumigatus* (57.9%) (Fig. 2A), with substantial numbers of gene families identified as accessory (28.3%) or singletons (13.8%). Accessory genes were more evenly distributed across the pan-genome (mean 1163 accessory gene families per genome, sd = 0.44) than singletons (mean 8 per genome, sd = 12.92) where high overall numbers of singletons were driven by a relatively small number of strains with high gene diversity (Fig. 2B-D). The distribution of accessory and singleton gene family abundance was not structured by phylogeny (Fig. 3), and the abundance of accessory and singleton genes was not significantly different between Clades. However, both accessory and singleton gene families were significantly related to genome size, with larger genomes associated with higher numbers of both accessory and singleton gene families (Fig. S5). Open pan-genomes are expected to accumulate genetic diversity in relation to the number of strains under evaluation, reflected in a higher number of accessory and singleton gene families relative to core gene families. Here, gene accumulation curves demonstrated a relatively closed pan-genome structure when considering only accessory gene families, but an open pan-genome structure with unsaturated genetic diversity when singletons were included (Fig. 2C). Although singleton gene families may represent overly stringent-binning of accessory gene families, the similar prevalence of singleton gene families between OrthoFinder and PIRATE defined homologues, and the relatively high number of singleton gene families with assigned InterPro Domains, leads us to conclude that these families may represent substantial genetic diversity in gene families that occur rarely in the greater population. The diversity of singleton genes in *A. fumigatus* suggests both a mechanism for the generation of substantial genetic diversity, and a mechanism for the purging of much of this diversity before these novel singleton genes become fixed in the population.

In fungi, novel genes are acquired either by horizontal gene transfer (HGT) (Friesen *et al*., 2006), functionalization via recombination, or duplication, and subsequent differentiation (Dutheil *et al*., 2016). Pan-genome size is influenced by population size, outcrossing frequency, and niche specificity (McInerney *et al*., 2017; Badet & Croll, 2020), where species with large populations, frequent gene flow, and substrate heterogeneity are associated with larger pan-genomes. For example, *Zymoseptoria tritici* is a highly outcrossing, widely distributed wheat pathogen with an estimated ratio of 60:40 core:accessory (Badet *et al*., 2020). For comparison, *Saccharomyces cerevisiae*, which although also widely distributed, but rarely outcrossing, has a ratio of 93.4:6.6 core:accessory (Li *et al*., 2019). Previous pan-genomic analysis of *A. fumigatus* have found ratios of core:accessory genes ranging from 83.29%:16.71% core:accessory using 12 isolates (McCarthy & Fitzpatrick, 2019), to 69%:31% core:accessory using 300 isolates (Barber *et al*., 2021). In this study, *A. fumigatus* has a ratio of approximately 67.2:32.8 core:accessory excluding singletons and 57.9:42.1 core:accessory including singletons. Ecologically, the exceptional diversity observed in the pan-genome of *A. fumigatus* is likely indicative of the massive population size and global distribution of the species, coupled to environmental heterogeneity and local adaptation to diverse niche opportunities. To tease apart the relative influence of each of these factors, boarder studies using the same methods across large datasets of fungal species with diverse distributions and lifestyle characteristics are needed.

While the vast majority of core gene families were assigned functional annotations (98.25%), this number was markedly lower for both accessory (79.48%) and singleton (81.27%) gene families. Given that core gene families are considerably more likely to be present in reference strains, and therefore more likely to be characterized, the high annotation frequency in core genes is expected. Conversely, accessory and singleton genes families had comparable frequencies of genes that could not be assigned to function. The complete absence of functional annotation for rare genes demonstrates the vast genetic diversity found in *A. fumigatus* and highlights the short-comings of single-reference based approaches for investigating population-wide genetic diversity.

We searched for signatures of clade-specific genetic diversity, and identified clade-specific gene families in all three clades (Fig. 4A). While the total number of unique gene families in each clade were intrinsically influenced by differences in clade size, we can generalize about the distribution of these gene families within clades. For example, while many clade-specific gene families were present in only a minority of isolates, others were present in all or nearly all of the isolates in a given clade. Presence/absence analysis also identified gene families that were uniquely absent among a single clade (Fig. 4B), including multiple clade-defining absences, where gene families were missing in all isolates of that clade, but present in nearly all isolates of the other two clades. Here, absence could represent either gene loss in the clade of interest, or gene gains in the other two clades. Overall, fewer absences were identified in Clade 1 than for the other two clades, with no gene families making our cutoff to define the loss as a Clade-defining feature. Conversely, clade-defining absences were identified in both Clade 2 (n=2 absences) and Clade 3 (n=23 absences).

Although infrequent, sexual recombination has been documented to occur in *A. fumigatus* (O’Gorman *et al*., 2009). However, the frequency of these recombination events in natural populations is unknown. Here, we characterized MAT-type across all isolates to look at the ratio of MAT1-1 to MAT1-2, where ratios close to 1:1 are indicative of random mating (Milgroom, 1996). Here, the presence of both mating types in all three clades indicates that the capacity for recombination exists in all clades. MAT loci proportions displayed a slight skew toward the MAT1-1 idiomorph both over all, and for each of the three clades with an overall MAT1-1:MAT1-2 ratio of approximately 7:5. The 11 strains with full length alignments to both MAT1-1 and MAT1-2 displayed different sequencing depth profiles for each idiomorph in all but three cases (AF100-1_3, IFM_59359, and IFM_61407) (Fig. S7), suggesting the potential for different underlying causes of the double mapping to both MAT loci in these strains. Interestingly, one of these strains Afu_343_P_11, was previously identified as an outlier (Abdolrasouli *et al*., 2015), for its exceptional genetic diversity compared with other strains from the same location. K-mer ploidy analysis supported haploidy in seven of the nine strains, but diploidy for AF100-1_3 and IFM_59359. Allele frequency analysis was in agreement with the depth analysis, supporting diploidy for AF100-1_3, IFM_59359, and IFM_61407, and haploidy for all others. Taken together, while low levels of diploidy may be present in our dataset, at least six of the nine strains with double MAT alignments may be more likely to represent low levels of noise (culture contamination, or tag switching during the sequence process). The diploid state is a signature of parasexual recombination and low incidence of *A. fumigatus* diploid strains have been previously found in *A. fumigatus* lung isolates form Cystic Fibrosis patients (Engel *et al*., 2020). While AF100-1_3 is a clinical Cystic Fibrosis lung isolate (Ross *et al*., 2021), IFM_59359 is a clinical isolate taken from a patient with pulmonary aspergilloma, and IFM_61407 is a clinical isolate from a patient with chronic necrotizing pulmonary aspergillosis (Takahashi-Nakaguchi *et al*., 2015). Thus, all three of these strains had the potential to be in the human lung environment for a long period of time.

Core gene families were primarily enriched for gene ontology terms related to essential functions such as transcription, translation, and transport. Accessory and singleton gene families were enriched for multiple terms related to metabolism-particularly terms related to nitrogen, carbohydrate, and phosphorus processing (Fig. 5A-C). Enrichment of clade-specific gene families echoed these results, with various metabolic processes enriched in clade-specific gene families in Clades 1 and 3 (Fig. 5D-E). To further investigate the potential for differential metabolic capacity across the phylogeny, we annotated CAZyme genes across all isolates to look for differences in CAZyme gene frequency between clades, and identified differential abundance in 28 CAZyme families (Fig. 6A), representing Auxiliary Activities, Carbohydrate-Binding Modules, Carbohydrate Esterases, Glycoside Hydrolases, and Glycosyl Transferases. These genes play a variety of roles in colonization, nutrient utilization, and substrate specificity (Zhao *et al*., 2013). Clade-specific differences in presence-absence and copy number of CAZymes may be driven by ecological and evolutionary factors relevant to environmental systems, such as niche occupation and subtle substrate specificity on various plant biomass materials. However, genetic differences driven by environmental ecological factors may also have clinical implications. For example, GH135 (Sph3, with an additional copy in Clade 3) is involved in the production of galactosaminogalactan (GAG), a critical component of *A. fumigatus* biofilms (Bamford *et al*., 2015). GH27 (with an additional copy in Clades 2 and 3) is structurally similar GH135, but currently uncharacterized in *A. fumigatus*. GH16 and GH55 (both with one less copy in Clade 3) are essential for proper *A. fumigatus* conidial morphogenesis (Mouyna *et al*., 2016; Millet *et al*., 2019). AA3 (with an additional copy in Clade 2, and AA7 (with one less copy in Clade 2, and two less copies in Clade 3) likely serve as oxidases with a role in the production of H_2_O_2_ for lignin depolymerization (Nekiunaite *et al*., 2016). Differences in the production of H_2_O_2_ may require corresponding differences in detoxification ability, and could impact interactions with host leukocytes that employ oxidative antifungal killing mechanisms (Morgenstern *et al*., 1997; Ibrahim-Granet *et al*., 2003; Philippe *et al*., 2003; Shlezinger *et al*., 2017; Santos *et al*., 2018). Metabolic specificity between isolates may also represent the possibility for niche preadaptation in clinical settings, as nutritional landscapes may not be uniform in clinical environments. For example, the cystic fibrosis lung environment is characterized by the impaired clearance and increased viscosity of mucins (Riordan *et al*., 1989). These mucins represent unique carbon and nitrogen sources which act as substrates for diverse microbial communities (Flynn *et al*., 2016) with the potential for both direct and syntrophic interactions between colonizing fungi and bacteria. To look for evidence of substrate specificity between the clades we used gene family homology to Af293 genes to identify clade-specific patterns in gene presence-absence variation in genes associated with nitrogen, carbohydrate, and phosphorus metabolism and found significantly differentially abundant genes in all three categories structured by clade (Fig. S10A-C). For example, Afu6g14620, which has been largely lost in Clade 2, but not Clades 1 and 3, encodes a putative Alpha-L-arabinofuranosidase, an enzyme which hydrolyzes arabinose side chains, and is known to be substrate specific (de Vries & Visser, 2001). Genes involved in secondary metabolism may also underlie clade specific niche occupation, as the expression of biosynthetic clusters is tied to differential nutrient access and nutrient sensing (Hewage *et al*., 2014). Here, we found substantial presence/absence variation in BGCs encoding diverse secondary metabolites (Fig. S12). Differential nutrient response between isolates, structured by gene variance or presence-absence variation in these gene clusters, may also have human disease implications. For example, the genes Afu3g13730, Afu3g13720, and Afu3g13710, which are lost in Clade 3 but present in nearly all other isolates are part of the same uncharacterized biosynthetic NRPS-like gene cluster (Bignell *et al*., 2016). Also coded under the GO terms for organonitrogen and organophosphorus metabolism, this cluster is preferentially expressed during the initial 4 hours of infection *in-vivo* (Bignell *et al*., 2016). Although the results presented here identify an intriguing direction for future study, functional work is needed to clarify realized differences in substrate usage, and any potential links between ecological niche occupation and realized differences in pathogenesis and virulence traits.

Another example of the potential pathogenesis and virulence impacts of gene variation is the discovery of loss of part of the gliotoxin biosynthesis gene cluster (Fig. S12). Gliotoxin is a powerful mycotoxin and an infamous virulence factor in *A. fumigatus* infection (Kwon-Chung & Sugui, 2009). A large deletion in eight strains, covering eight genes on Chromosome 6 (relative to Af293), included the tailoring and structural genes *gliI* and *gliJ,* the transcription factor *gliZ,* and a partial deletion in the NRPS gene *gliP*. Among these, *gliI* and *gliP* are considered essential to gliotoxin biosynthesis (Cramer *et al*., 2006; Scharf *et al*., 2012). Whereas the full *gliP* protein contains two sets of canonical NRPS A-T-C modules plus a tailing T (thioesterase), the truncated gene is predicted to encode a single A-T-C module and a tailing A, making peptide formation unlikely (Fig. S13). As the production of gliotoxin is thought to impart enhanced survival and infection persistence (Bruns *et al*., 2010), the presence of this large deletion in eight strains representing a persistent lineage present in a Cystic Fibrosis patient with long-term Aspergillosis (Ross *et al*., 2021), hints that gliotoxin production is not necessary for persistence in patients with cystic fibrosis, and highlights possible alternative/adaptive roles for the functional loss of this cluster under some conditions.

To test if clinically-relevant alleles were also biased in their distribution across the three clades, we conducted variant scans on a set of 13 genes associated with antifungal drug resistance. These included targeted analysis of six functionally characterized amino acid changes available for six genes (*cyp51A*, *cyp51B*, *hapE*, *hmg1*, *cox10* and *fks1*) (Table S2), as well as surveying all amino-acid changing variants across these six genes and an additional seven genes associated with azole resistance, but lacking functional characterization of specific amino acid changes. Targeted scans only identified previously characterized mutations only in the *cyp51A* gene (Fig. 6B). While the azole resistant TR34/Leu98His and Gly138Cys genotypes were found exclusively in Clade 2, other characterized variants in *cyp51A*-mediated azole resistance were scattered in low abundance throughout Clade 1 (Pro216Leu, Met220Val) or Clade 2 (Gly448Ser, His147Tyr), or represented in both Clade 1 and Clade 2 (Gly54Glu). Conversely, no characterized *cyp51A* resistance variants were found in Clade 3. An additional scan for all non-synonymous variants in *cyp51A* found 11 amino acid changes that occurred frequently across the phylogeny. Interestingly, two of these changes (Glu255Asp and Thr248Asn), are variants that map to a monophyletic branch containing the Af293 reference Clade 1 isolates with signs of introgression, making the Glu255Asp and Thr248Asn genotypes the rule rather than the exception, and highlighting the potential problems associated with defining mutations relative to a single reference. Similarly, three variants (Lys427Glu, Val172Met, and Tyr46Phe), were absent only from the monophyletic branch containing the reference and from all Clade 3 isolates. Finally, six additional variants were present only in low abundance and exclusive to Clade 1 (Ala9Thr, Lys427Arg, Ile242Val, Ala284Thr), or Clade 2 (Ser297Thr, Phe495Ile). While the functional impacts of these low-abundance amino acid changes are unknown, the frequency of uncharacterized *cyp51A* amino acid changes deserves future consideration. Additionally, while the genes scanned did not contain characterized drug resistance alleles, they all contained multiple uncharacterized amino acid changing variants (Fig. S11). Several of these changes demonstrated phylogenetic structure, such as the Leu76Phe change in AFUA7G01960, which was both prevalent and nearly exclusive to Clade 2 isolates. *cyp51A* mutations are the most well studied resistance mechanisms in *A. fumigatus* but only account for an estimated 43% of resistant isolates (Bueid *et al*., 2010). The distribution of nonsynonymous variants in other genes associated with resistance highlight the importance of future research on the functional impact and population distribution of alternative drug resistance mechanisms in *A. fumigatus*.

The availability and continued improvement of high quality reference genomes has enabled insights into the evolution and variation of fungal genome structure (Hartmann *et al*., 2017), intragenomic evolutionary rates (Croll *et al*., 2015; Laurent *et al*., 2018), and how intraspecific genetic variability is structured across fungal populations (Wyss *et al*., 2016; Cissé *et al*., 2018). However, there is growing appreciation that a single reference genome is incapable of capturing the genetic variation present across a species. Whereas genes absent in the reference genome are likely to be ignored using reference based alignment, *de novo* strategies for genome assembly offer the possibility of capturing the full repertoire of genetic diversity (Potgieter *et al*., 2020). Here, we used a combined genomics approach to leverage both reference-based and *de novo* strategies to illuminate population structure and genetic diversity across the *A. fumigatus* pan-genome. These populations, subdivided into three primary clades, are characterized by exceptionally high gene diversity and substantial presence/absence variation, representing one of the largest fungal-pan genomes ever reported. While our ancestry and LD analysis suggest that recombination occurs at high rates within-clades than between clades, laboratory studies are needed to confirm the recombination rate, frequency of mating, and mating compatibility within and between isolates from the three primary lineages. The genes defining the thee primary clades are enriched for diverse metabolic functions, hinting that population structure may be shaped by environmental niche occupation or substrate specificity. If niche occupation translates to realized differences in nutrient usage or stress tolerance, it may have disease initiation and/or progression implications (Cramer & Kowalski, 2021). Finally, as evidenced by the numerous gene families identified here which have no homologue in Af293, the under-characterization and inability to assign annotations to approximately 20% of the accessory genome, and variant scans which identified clear cases where Af293 was the exception, rather than the rule for the population (and therefore inappropriate to define mutations against). This work highlights the inadequacy of using single reference-based approaches to capture the genetic variation across a species, and the power of combined genomics approaches to elucidate intraspecific diversity. As this study can only represent conclusions based on the data available, we anticipate that as additional genome sequences become available, novel genomic and phylogenetic diversity will continue to be discovered in *A. fumigatus*. Additionally, the methodological hurdles associated with assessing population structure and recombination frequency in organisms with complex clonal-sexual life cycles is significant and highlights the need for the further development of computational tools capable of addressing issues specific to fungi.

## ACKNOWLEDGEMENTS

LAL, JES, and RAC are supported by funding from the National Institutes of Health (grant no. R01AI130128), to RAC, with additional support to LAL and JES from a UC Riverside-City of Hope seed grant to JES. JES is a CIFAR Fellow in the program Fungal Kingdom: Threats and Opportunities and was supported by funding from National Science Foundation (grants no. DEB-1441715 and DEB-1557110). All analyses and data management was performed on the High-Performance Computing Center at the University of California, Riverside, in the Institute of Integrative Genome Biology, supported by grants from NSF (DBI-1429826) and NIH (S10-OD016290). The authors would like to thank Dr. Tobias Hohl, Dr. Michail Lionakis, Dr. Sean Lockhart, Dr. John Wingard, and Dr. Alix Ashare for providing *A. fumigatus* isolates. Special thanks to Ben Auxier for his comments on a previous version of this manuscript.

## DATA AVAILABILITY

All newly sequenced genomes have been deposited into GenBank: accession numbers for each sequencing project can be found in Table S1. All programing scripts associated with this project are available at: https://github.com/MycoPunk/Afum_PopPan. The definitive version of the pipeline used for VCF generation can be accessed at https://github.com/stajichlab/PopGenomics_Afumigatus_Global. All data associated with this project are available from DOI: 10.5281/zenodo.5775265

## AUTHORS’ CONTRIBUTIONS

LAL, JES and RAC planned the project. LAL wrote the manuscript with editorial input from JES and RAC. RAC sequenced genomes. BSR performed DNA extractions on *A. fumigatus* strains sequenced in this project. JES carried out assembly and annotation. LAL and JES carried out population genomics, phylogenomics, and pan-genomics.

## FIGURE AND TABLE LEGENDS

**Fig. S1: fastSTRUCTURE marginal likelihood values.** Marginal likelihood increased until K=5, with the largest increase occurring between K=1-3.

**Fig. S2: Population sub-structure.** Population substructure within the three clades, was investigated using iterative DAPC and fastSTRUCTURE analysis of vcf SNP sites subset by clade. **A)** The optimal number of PCs for the sub-clades was determined to be 5 for the substructure within Clade 1. **B)** The optimal number of PCs for the sub-clades was determined to be 3 for the substructure within Clade 2. **C)** The optimal number of PCs for Clade 3 was determined to be 1, and was therefore not further analyzed with DAPC. **D)** Iterative DAPC analysis supported 5 sub-clades within Clade 1. **E)** Iterative DAPC analysis supported 5 sub-clades within Clade 2. **F)** Iterative fastSTRUCTURE at K=5 supported frequent introgression between sub-clades in Clade 1 (87 out of 200 strains) **G)** Iterative fastSTRUCTURE at K=5 supported frequent introgression between sub-clades in Clade 1 (7 out of 45 strains). Sub-clade designations were not consistent between DAPC and fastSTRCTURE (i.e. DAPC sub-clade1.1 is not equivalent to fastSTRUCTURE sub-clade1.1).

**Fig. S3: The Influence of sample size on LD-decay.** To investigate the influence of sample size on LD-decay, we iteratively sampled n = 5, 10, 20, 50 or 100 isolates without regard to population structure (each n averaged over 20 independent iterations), and calculated LD50 in base pairs (BP). The influence of sample size on LD estimates showed a strong inverse relationship between sample size and LD decay, with an LD50 of 1,489.55 BP at n=5, 308.80 BP at n=10, 5.02 BP at n=20, 4.7 BP at n=50, and 2.76 BP at n=100. **A)** linear-linear plot. **B)** zoomed in log-linear plot, with arrows denoting LD50 at each n.

**Fig. S4: Pan-genome gene family distribution using PIRATE. A)** The pan-genome of 260 *A. fumigatus* strains included 15,476 gene families in total, including 8,600 (55.57%) core genes (present in >95% of strains), 3,618 (13.92%) accessory genes (present in >2 and < 248 strains) and 3,258 (21.05%) singletons (present in only one isolate). **B)** The distribution of the number of genomes represented in each gene family. **C)** Gene family accumulation curves, including (green) and excluding (yellow) singletons. **D)** The distribution of unique accessory gene families by strain. **E)** The distribution of unique singleton gene families by strain.

**Fig. S5: Relationship between genome size and number of gene families.** Both **A)** accessory gene families and **B)** singleton gene families were significantly related to predicted genome size, although this relationship yielded low R^2^ values, particularly for singleton gene families.

**Fig. S6: Proportion of Secreted Proteins across the pan-genome by frequency and clade.** Secreted Proteins were predicted using the programs Signal P and Phobius. In most cases, Phobius predicted a slightly greater number of secreted proteins than Signal P. Both Signal P (lightest shading) and Phobius (mid-level shading) contained proteins not annotated by the other program, but most individual proteins called as secreted overlapped between the two prediction programs (darkest shading) **A)** Proportion of gene families predicted to represent secreted proteins out of all gene families by frequency (core, accessory, and singleton) and **B)** by Clade for clade-specific gene families. In Clade 3, only two categories are depicted as all secreted proteins predicted by Signal P were also predicted by Phobius. Significant differences were assessed using Pairwise proportion tests with Bonferroni adjustment for multiple comparisons at *p* < 0.05. Letters (lowercase) in common, indicate no significant difference between groups. Significance results were consistent across Signal P, Phobius and Consensus inferred secreted proteins.

**Fig. S7: Depth profiles of MAT idiomorph alignments** for the nine isolates with significant alignments to both MAT1-1 and MAT1-2, plus positive controls. All strains had alignments over the entirety of the MAT region for both idiomorphs, but these alignments different in depth, suggesting different underlying explanations. For MAT1-2 alignments, the trailing ∼270 bp region that is conserved between MAT1-1 and MAT1-2, is visible here in the control as well as in several of the isolates that mapped to both idiomorphs. While MAT1-2 idiomorph strains have a full alignment over the (∼1078bp) MAT1-2 reference, MAT1-1 idiomorph strains only align over the ∼270bp conserved region of the MAT1-2 reference.

**Fig. S8: K-mer profiles of strains with alignments to both MAT1-1 and MAT1-2** while seven of the nine strains display single peaks indicative of haploidy, two strains, AF100-1_3 and IFM_59359 display low secondary peaks, potentially indicative of diploidy.

**Fig. S9: Allele frequency profiles of strains with alignments to both MAT1-1 and MAT-2** Plots indicate genome-wide allelic ratios across all SNPs for each genome, where 1 or 0 indicates homozygosity (1 indicating a polymorphism, and 0 matching the reference allele of Af293). Although homozygous positions are expected to dominate the data (outer graph), diploids are expected to have an additional peak at an allele frequency of 1/2, visible when 1 and 0 are removed from the graph space (inner plot). While six of the nine strains display no notable peaks at 1/2, three strains (AF100-1_3, IFM_59359, and IFM_61407) do show peaks at 1/2. Four other strains, while likely haploid, display significant areas of copy number variation as evidenced by additional peaks just below 1/4 and just above 3/4.

**Fig. S10: Presence/absence variation in metabolic genes.** Gene family clusters with significant homology to annotated genes the Af293 reference genome were assigned using BLASTP (e-value < 1e^-15^). Using these annotated clusters, we identified differential abundance in clusters with Af293 annotations between the three clades, for Af293 genes in FungiDB under the GO terms **A)** organonitrogen compound metabolic process (GO: 1901564, n = 25 significantly differentially abundant genes. Only the first seven are depicted for visualization), **B)** carbohydrate metabolic process (GO: 0005975, seven significantly differentially abundant genes), and **C)** organophosphate biosynthetic process (GO: 0090407, five significantly differentially abundant genes). Genes denoted with * are shown grouped with GO: 1901564, but are also annotated in GO: 0090407.

**Fig. S11: Distribution of non-synonymous variants in non-*cyp51A* genes associated with fungal drug resistance.** In addition of *cyp51A*, a database of other genes known to be associated with antifungal drug resistance was assembled (Table S2) and all isolates were scanned for non-synonymous variants in these genes. Four genes, Cyp51b, cox10, mdr2, and AFUA 7G01960 contained uncharacterized non-synonymous amino acid changes, several of which demonstrated phylogenetic structure.

**Fig. S12: Presence/absence variation in secondary metabolite biosynthetic gene clusters.** Gene family clusters with significant homology to annotated genes the Af293 reference genome were assigned using BLASTP (e-value < 1e^-15^) and genes in gene clusters encoding notable *A. fumigatus* secondary metabolites (as defined in (Bignell *et al*., 2016)) were mapped onto the phylogeny in R. Out of 230 genes in 26 clusters, seven were convoluted with one other gene in the cluster, where both genes clustered into the same gene family (in these cases both genes are listed separated by /). Three genes (Afu1g10275 in cluster 2, Afu7g00140 in cluster 23 and Afu8g00450 in cluster 25) could not be confidently assigned to an Orthofinder gene family and are not displayed. BGC = Biosynthetic gene cluster.

**Fig. S13: Presence/absence variation in gliotoxin cluster genes.** The boundaries of the deletion were the same for all eight strains and included the partial truncation of gliP, covering one A-T-C module.

**Table S1: Accession numbers and strain information.**

**Table S2: Database of characterized antifungal resistance mutations in *A. fumigatus*.** Database was assembled from the literature, and supplemented with search results from the MOADy antifungal drug resistance database.

**Table S3: DAPCA clade assignments, including posterior probabilities for each strain used in this study.**

**Table S4: fastSTRUCTRE clade assignments, including posterior probabilities for each strain used in this study.**

**Table 5: DAPC sub-clade assignments, including posterior probabilities for each strain.**

**Table S6: fastSTRUCTURE sub-clade assignments, including posterior probabilities for each strain.**

**Table S7: Abundance and annotation of clade-specific gene families.**

**Table S8: CAZyme gene counts for all A. fumigatus isolates.**

**Table S9: CAZyme enrichment stats table.** Mean gene counts and host-hoc results for the 28 CAZymes with significantly different abundance between the three Clades. Letters (in parenthesis) in common indicate no significant difference in abundance.

