## Supplementary figures and images for "Combined Pan-, Population-, and Phylo-Genomic Analysis of *Aspergillus fumigatus* Reveals Population Structure and Lineage-Specific Diversity"

### Supplemental Fig. S1

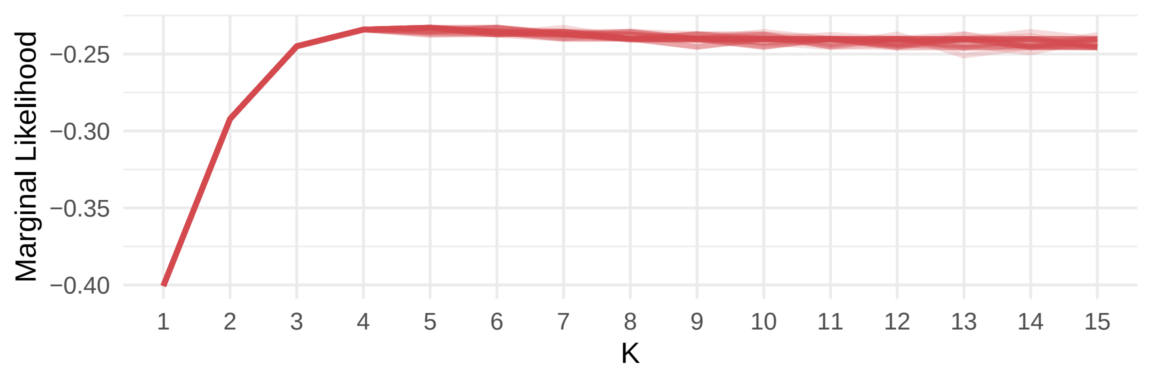

### Supplemental Fig. S2

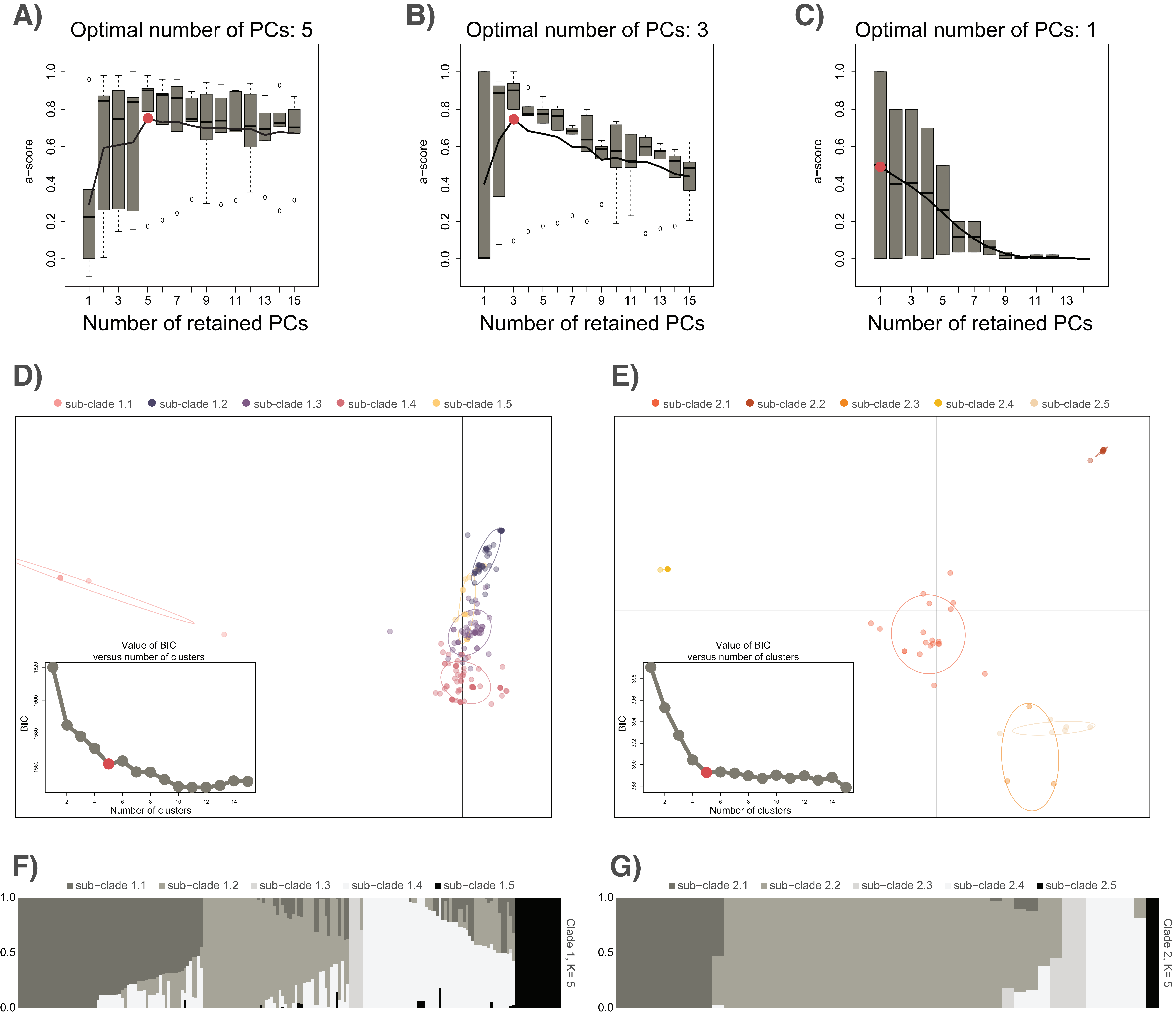

### Supplemental Fig. S3

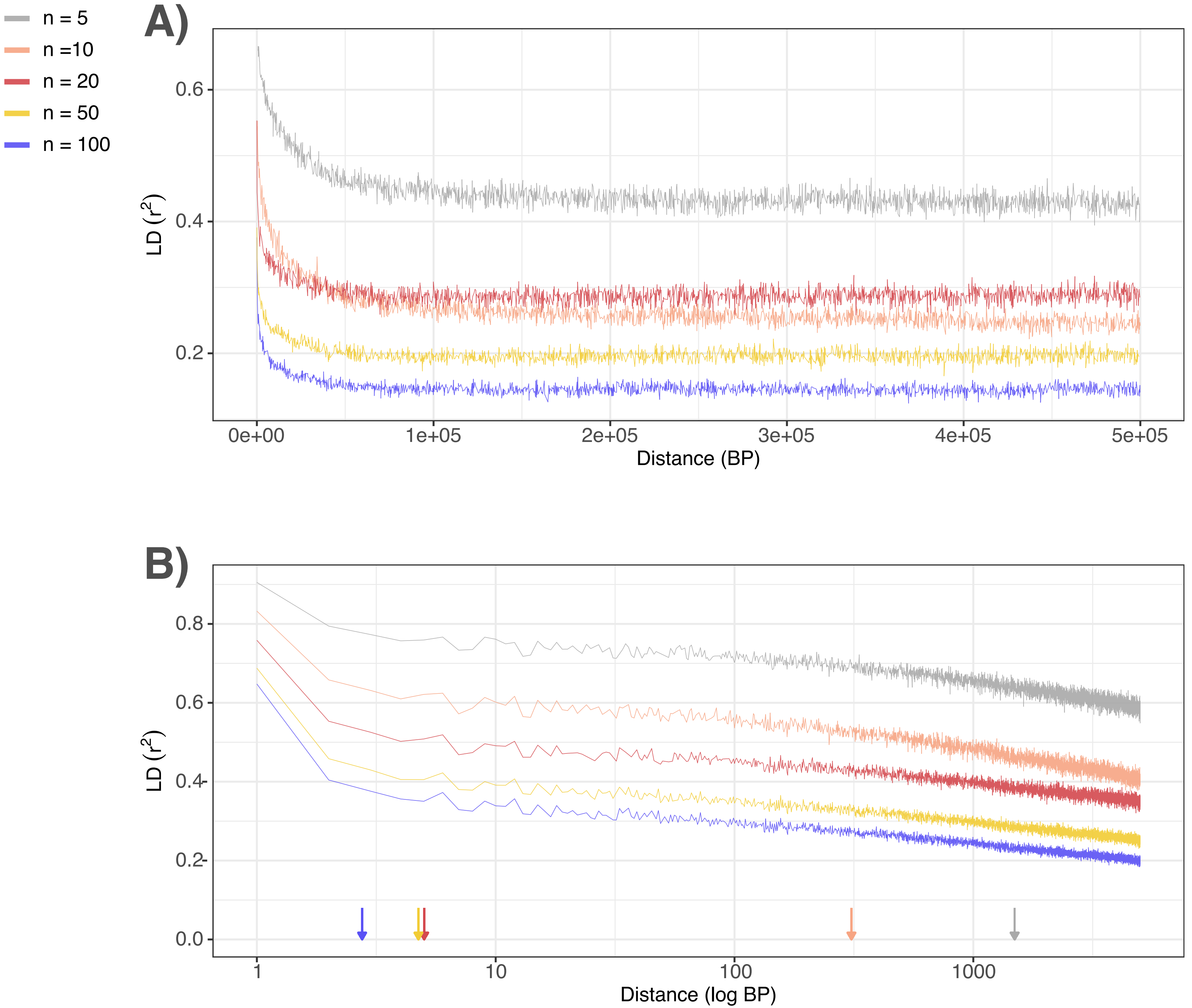

### Supplemental Fig. S4

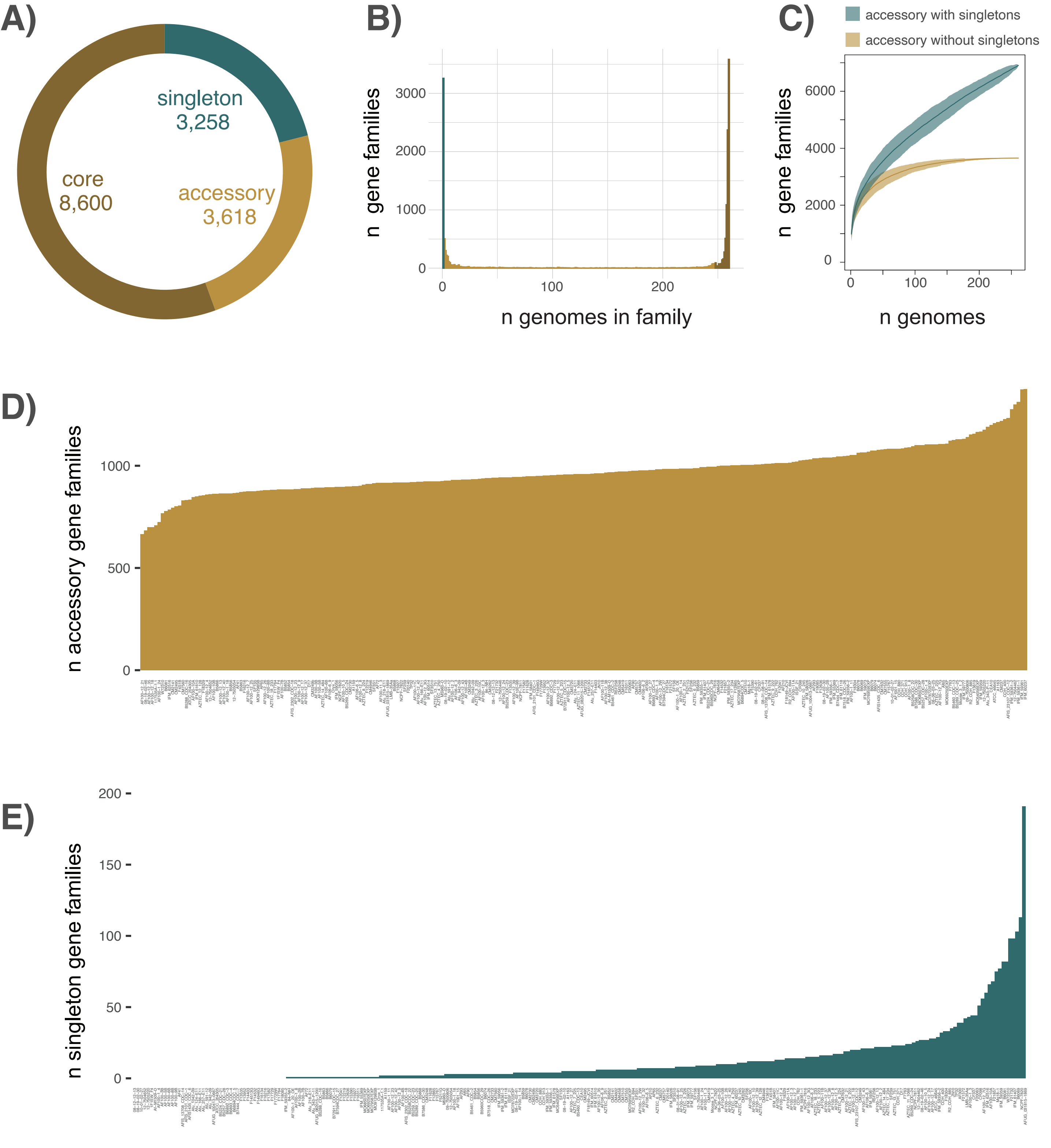

### Supplemental Fig. S5

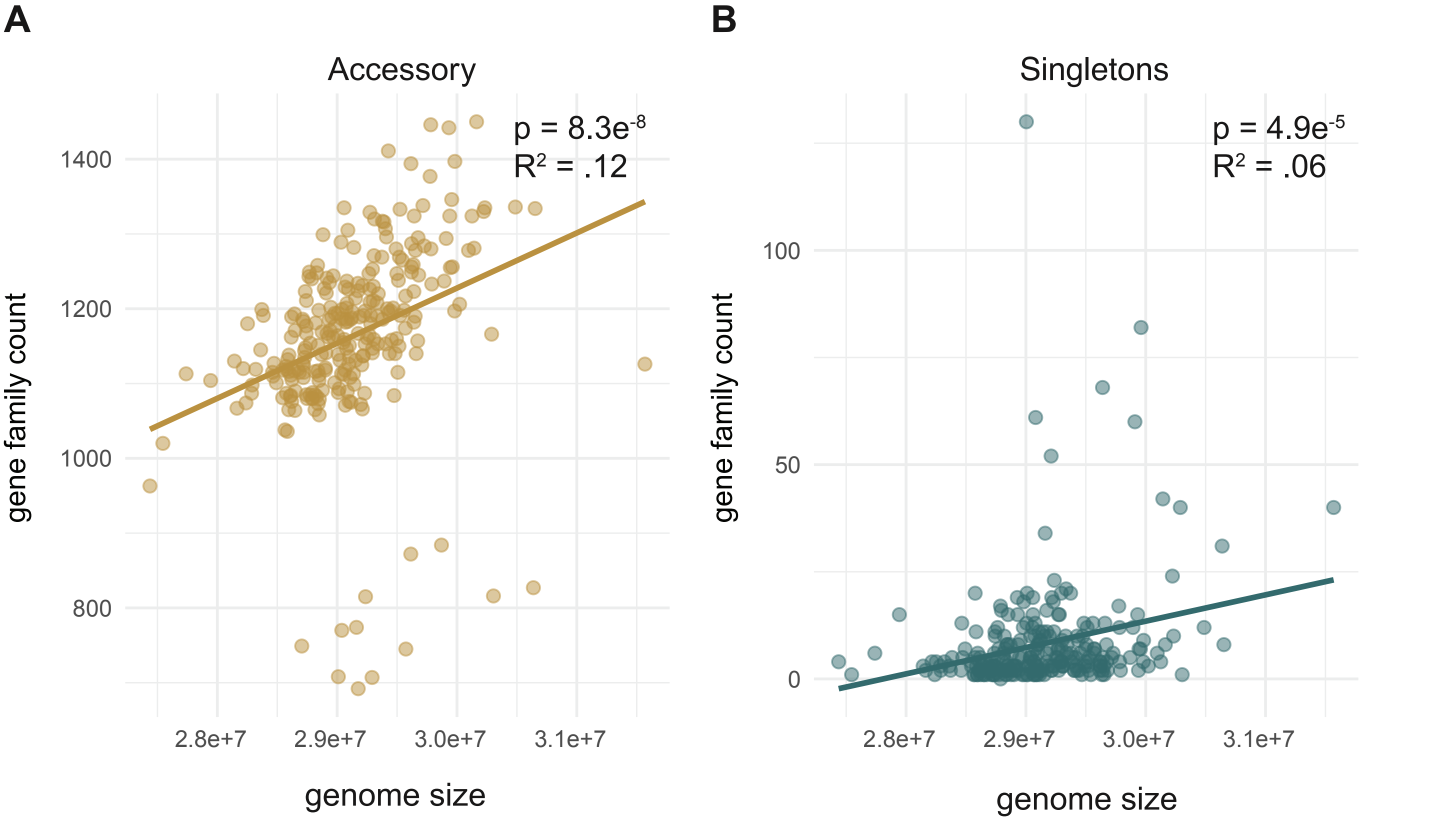

### Supplemental Fig. S6

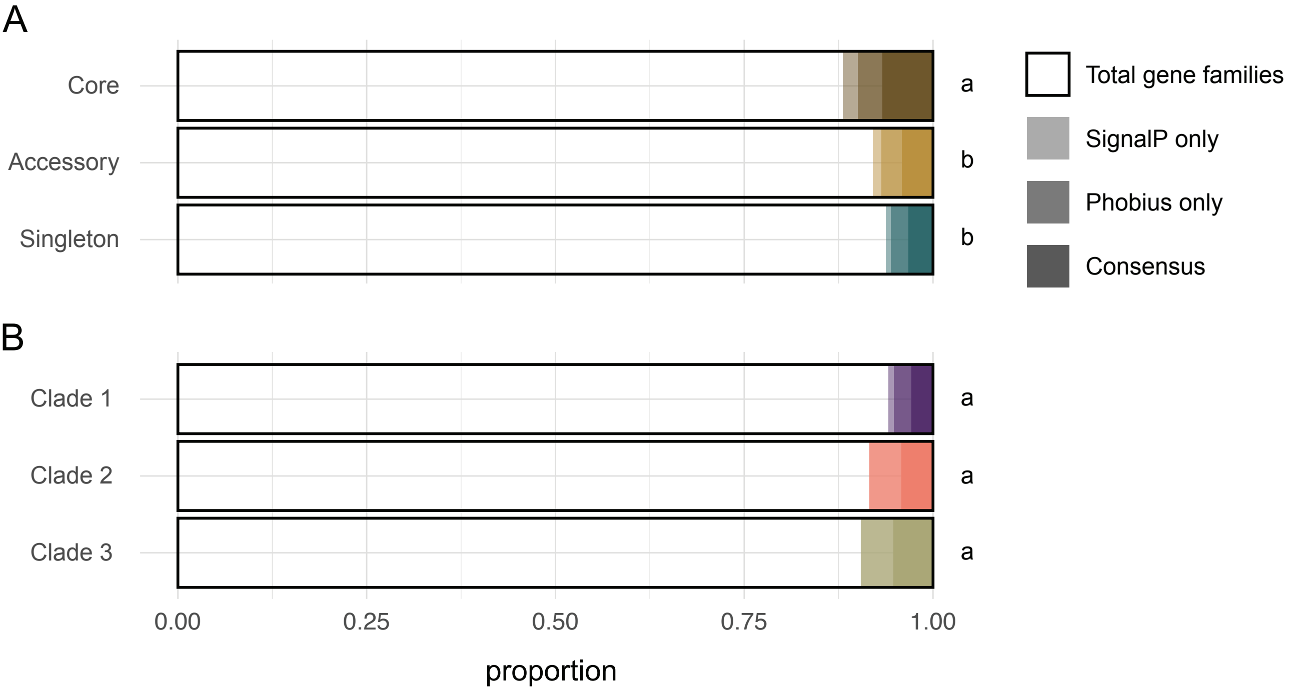

### Supplemental Fig. S7

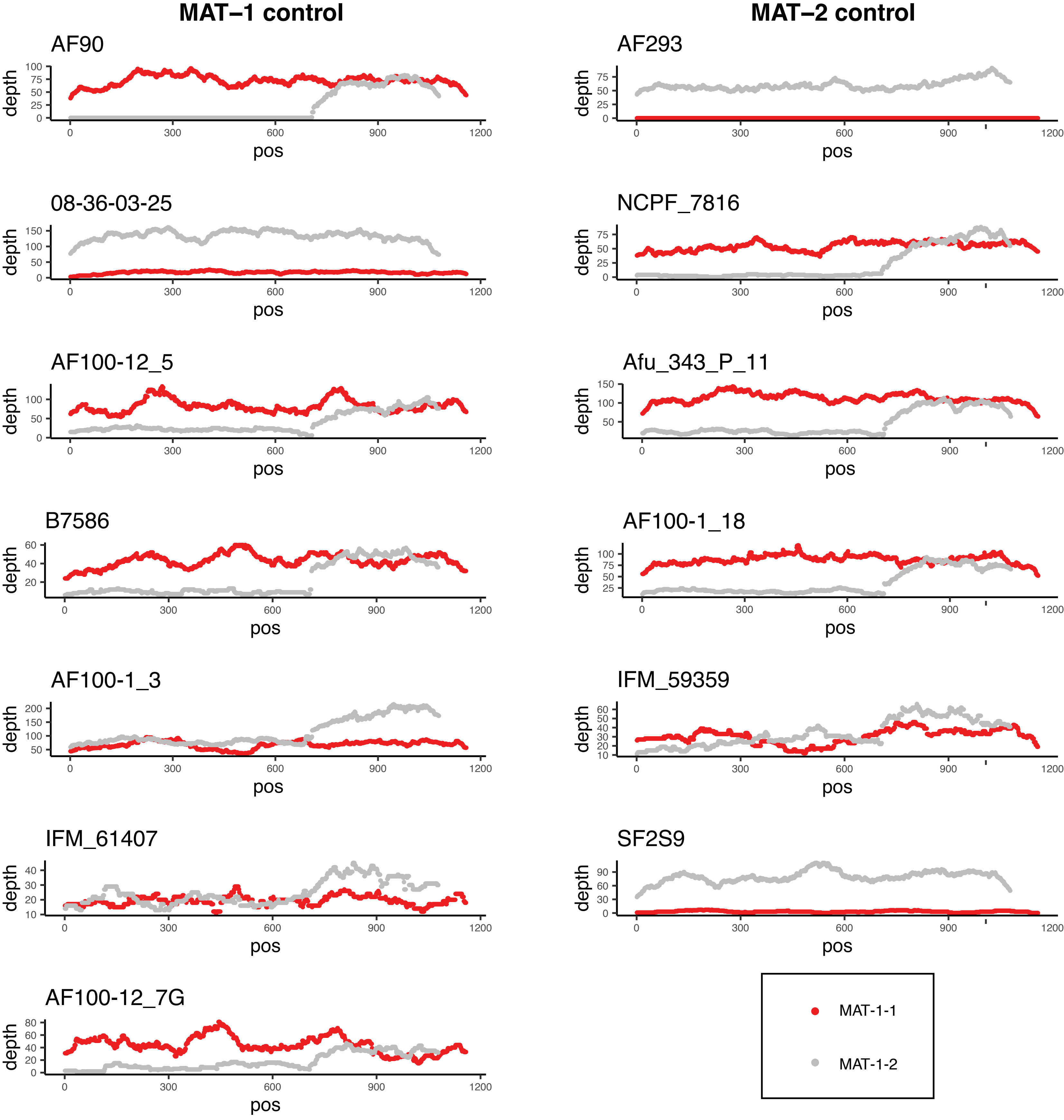

### Supplemental Fig. S8

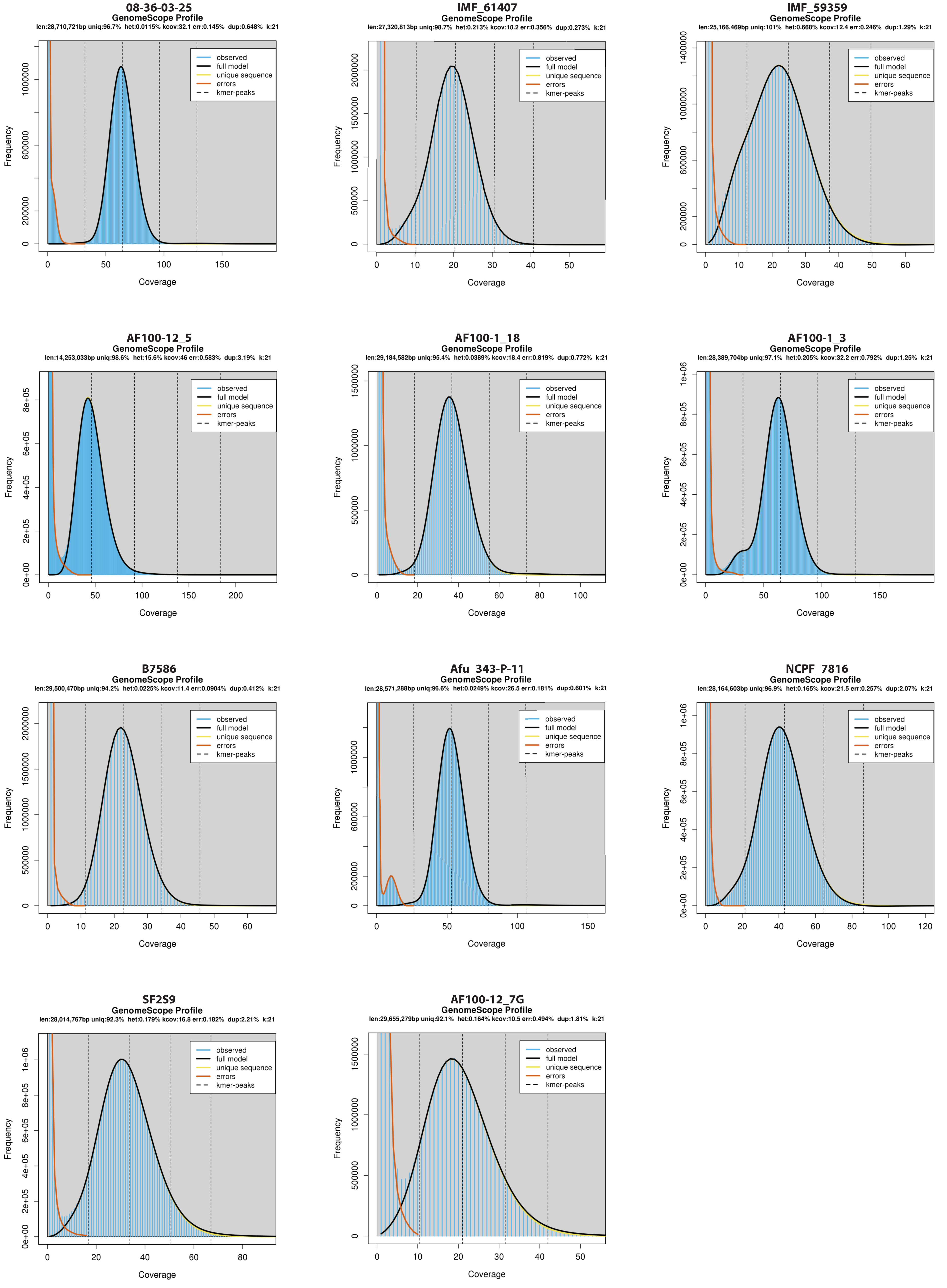

### Supplemental Fig. S9

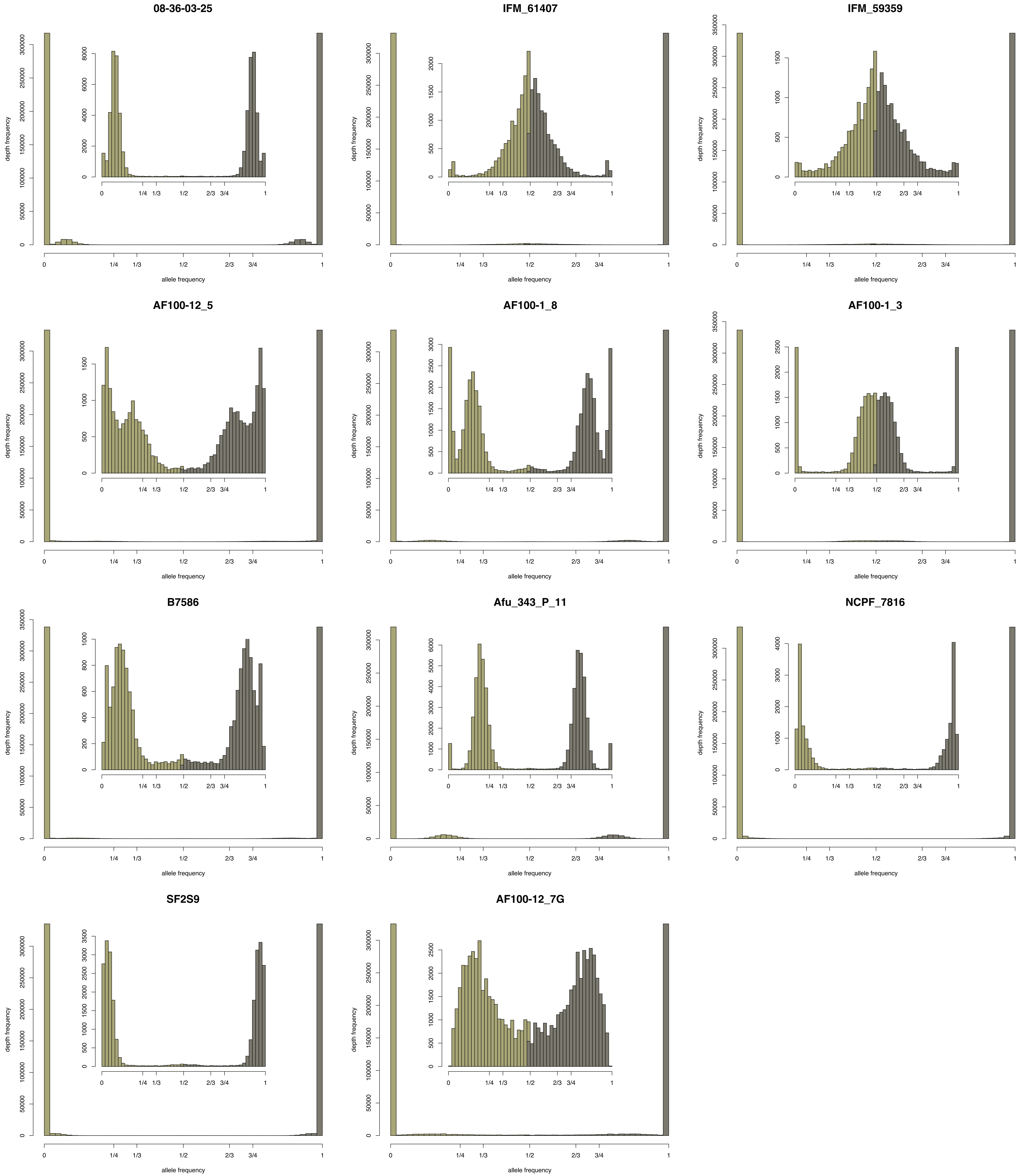

### Supplemental Fig. S10

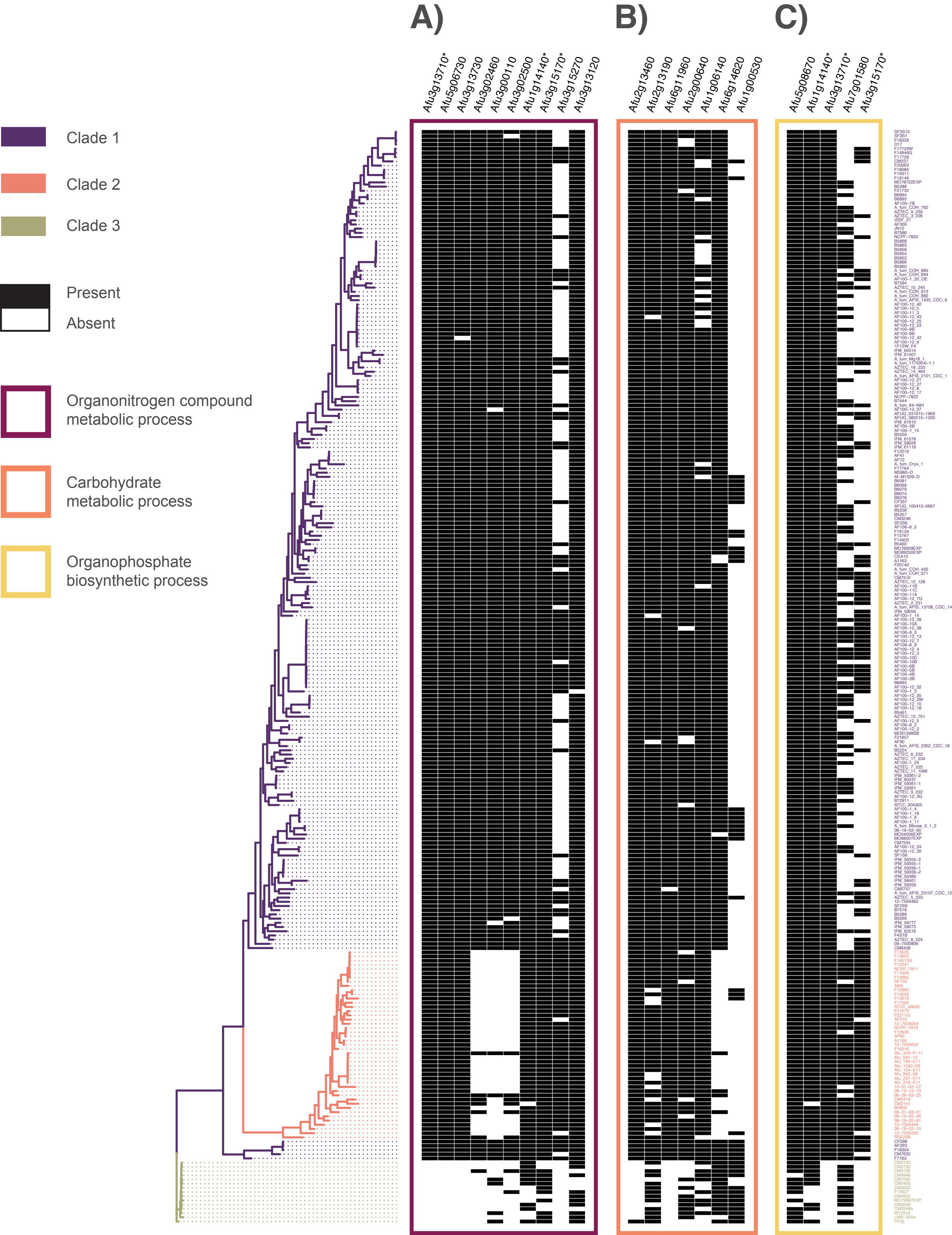

### Supplemental Fig. S11

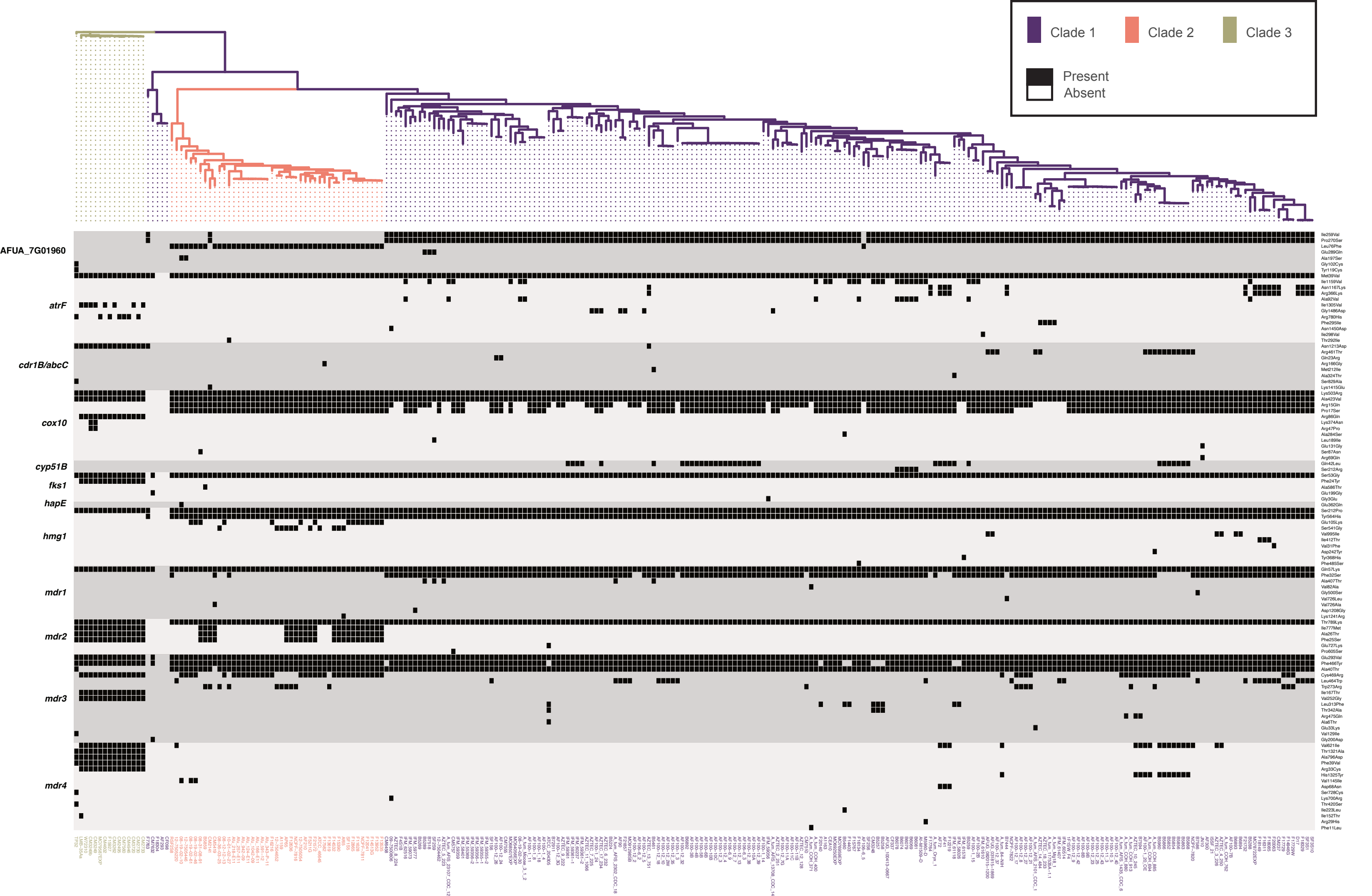

### Supplemental Fig. S12

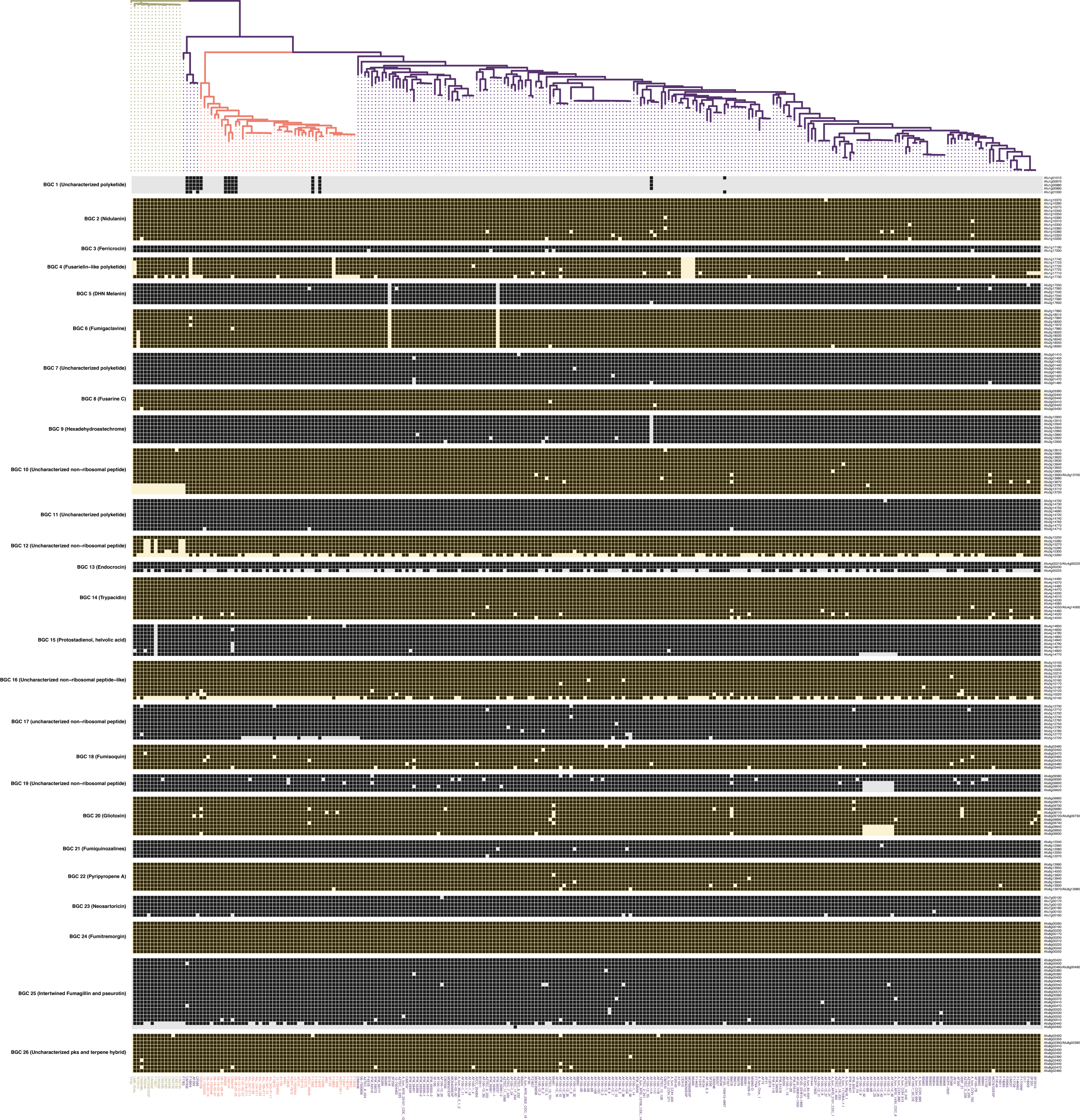
