## Supplemental Fig. S13 for "Combined Pan-, Population-, and Phylo-Genomic Analysis of *Aspergillus fumigatus* Reveals Population Structure and Lineage-Specific Diversity"

### Afu6g09660 (gliP)

Carrier = peptidyl carrier protein (PCP) or T site
gliP truncation architecture= one ATC module + A: A−T−C−A
gliP Af293 architecture= two ATC modules + T: A−T−C−A−T−C−T

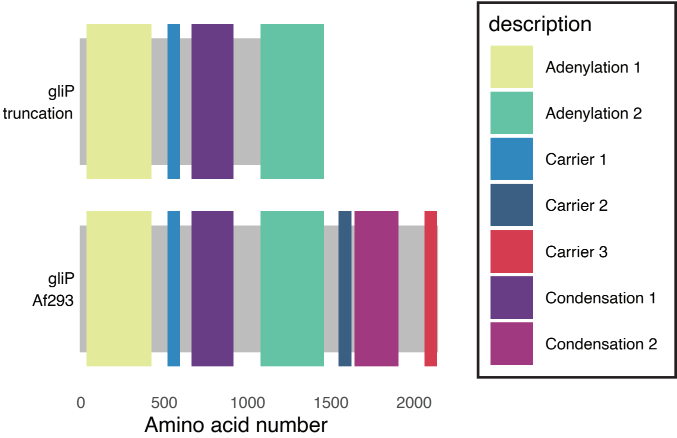
